# Adaptive immunity is dispensable for salamander appendage regeneration

**DOI:** 10.64898/2026.04.13.718117

**Authors:** Chimezie Harrison Umeano, Marlene Oesterle, Ameneh Ghaffarinia, Ugonna Okeke, Natalie Larsson, Sapna Meena, Emma Bolech, Camila R. Consiglio, Elif Eroglu, András Simon, Nicholas D. Leigh

## Abstract

Complex multi-tissue regeneration capacity varies across vertebrates. Mammals are amongst the least regenerative species, while salamanders can regenerate complex tissues such as limbs and tails throughout life. Previous studies have shown that innate and adaptive immune cells are present during salamander limb regeneration. While innate immune cells have been shown to promote limb regeneration, it is unknown whether adaptive immunity is responsive to amputation or plays a role in appendage regeneration. Here we show that during limb regeneration in axolotls, the immune response is characterized by a coordinated immunoregulatory signature including the downregulation of antigen presentation, cytokine secretion, and T cell activation. We corroborate this transcriptomic data *in vivo* using skin allografts in newts and define the blastema as an immunosuppressed niche. To test the role of adaptive immune cells in regeneration, we generated *Recombination activating gene 1* deficient (*Rag1*^-/-^) newts. *Rag1*^-/-^ newts lack antigen receptor recombination and show a marked reduction of adaptive immune cells. We find that *Rag1*^-/-^ newts do not reject allografts, confirming their functional immunodeficiency. Finally, we demonstrate that both larval and adult newts regenerate appendages in the absence of adaptive immunity. Our work demonstrates that the adaptive arm of the immune system is not required for appendage regeneration and establishes an important model for novel experimental approaches in comparative immunology and regenerative biology.

## Introduction

Regenerative capacity varies across species. Salamanders are the only tetrapods able to regenerate limbs and various other complex tissues throughout life^1–4^. Appendage regeneration is dependent on the blastema—a mass of lineage-restricted progenitor cells formed at the injury site which proliferates, differentiates, and reorganizes to replace the lost tissue^5^.

The formation of the blastema depends on a specialized epidermis (i.e., wound epidermis) and innervation^6–12^. More recently, phagocytes have also been demonstrated to be required for limb regeneration^13^. This requirement for phagocytes was confirmed in various species and models of complex tissue regeneration, including in mammals^14–17^. Though phagocytes have been shown to be required, we have limited knowledge of their functional contributions to regeneration.

A primary function of phagocytes is to activate and control adaptive immune responses, which are pivotal for pathogen clearance and immunological memory. Previous work has shown that, in addition to innate immune cells, there are adaptive lymphocytes in the regenerating limb^18,19^. Regarding a potential role for adaptive lymphocytes, studies have demonstrated that during appendage regeneration there is a decrease in circulating lymphocytes^20^, and treatment with immunosuppressants can impede appendage regeneration^21,22^. Additionally, newts with an ongoing allograft response prior to limb amputation display retarded regeneration^23^. These studies demonstrate correlative associations between amphibian appendage regeneration and adaptive immune responses, although direct interactions between these processes remains to be demonstrated.

Since salamanders maintain lifelong appendage regenerative capacity, it is also notable that adaptive immune function varies dramatically across life. Generally, the adaptive immune system of embryonic and larval/neonatal animals has a strong bias towards tolerogenic over inflammatory responses^24^. A tolerogenic bias has been shown to coincide with high regenerative capacity. For example, the regenerative neonatal mouse heart benefits from the immunosuppressive influences of neonatal macrophages, and B and T cells to facilitate regeneration^25,26^. As neonatal tolerance erodes during maturation, regeneration capacity is lost^25,26^. Similar shifts in baseline immune responsiveness occur in *Xenopus* and correlate with the loss of the ability to regenerate appendages^27^. How the developing and mature adaptive immune system of salamanders influences regeneration remains unresolved.

These changes in immune responsiveness are tied to metamorphosis—the transition from larval to adult form. Skin allograft rejection has served as an informative means to study amphibian immunity and the effects of metamorphosis on immune function^28^. In larval *Xenopus,* skin allografts are chronically rejected, while post-metamorphosis allograft responses become acute^29^, mirroring allograft responses of mammals. Salamanders diverge from this standard, as larval salamanders tolerate allografts, and adult salamanders reject allografts chronically^30^. This positions salamanders as an outlier regarding their adaptive immune function, though whether this deviant function across metamorphosis is a key to regeneration remains to be explored.

Salamanders are thus the only tetrapod capable of lifelong appendage regeneration and demonstrate unique adaptive immune function compared to their less regenerative amphibian counterparts (e.g., *Xenopus*). It is however unknown how the adaptive immune system interacts with regenerative tissue and more importantly whether adaptive immunity is required for regeneration.

Here, we show that after limb amputation in axolotl both innate and adaptive immune cells acquire transcriptional signatures consistent with immunosuppression compared to their counterparts in non-regenerating limbs. By skin allograft into the blastema of the newt, *Pleurodeles waltl*, we validated this transcriptomic data *in vivo*, demonstrating that the regenerative niche is immunosuppressed. To test for a role of adaptive immunity in regeneration, we generated *Recombination activating gene 1* deficient (*Rag1*^-/-^) newts which lack mature B and T cells. We show that this immunodeficiency results in the inability to reject allografts. Using these immunodeficient newts, we show that adaptive immunity is dispensable for larval and adult newt appendage regeneration. By generating an immunodeficient newt line, this study substantially advances our understanding of the interplay between immunity and appendage regeneration by demonstrating that regeneration proceeds unperturbed in the absence of mature adaptive immune cells. This line has broad implications for the regeneration field, now allowing for previously intractable allo- and potentially xenografts, by establishing a universal recipient system for regeneration studies. Altogether, these findings add a critical new layer of understanding to the required inputs for complex tissue regeneration, which will aid future efforts aimed at coaxing regeneration in mammals.

## Results

### Limb amputation and subsequent regeneration invoke a cellularly diverse innate and adaptive immune response

While the immune system is required for limb regeneration^13^, little is known about the immune cells present during regeneration and their contributions. Taking advantage of the high-quality axolotl genome assembly^31^, we analyzed single cell RNAseq (scRNAseq) data across a time course of limb regeneration^19^. To understand how immune cells were impacted during regeneration we compared non-regenerating limbs (both intact and contralateral to an amputation) to regenerating limbs (Figure 1A). Although contralateral limbs may be exposed to systemic signals associated with regeneration, they are not exposed to the local regenerative niche^32^. Comparisons between grouped non-regenerating controls and regenerating limbs were thus designed to enrich for immune adaptations specifically associated with local regenerative processes and not systemic immune responses.

**Figure 1:**
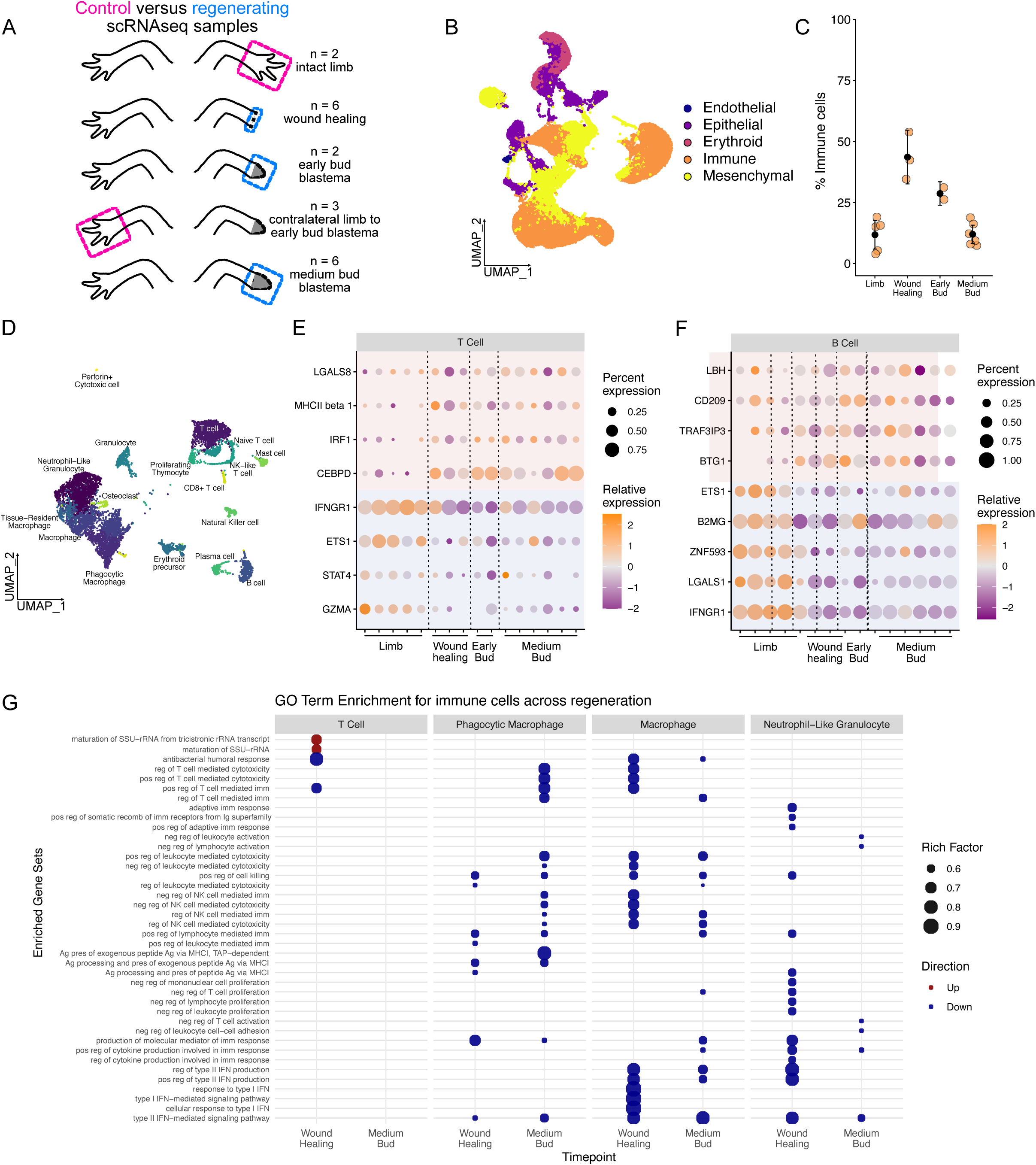
Limb amputation invokes a cellularly diverse immune response. **(A)** Samples analyzed using scRNAseq. Magenta signifies control samples which contain homeostatic limb and contralateral limbs while blue marks regenerating samples that were compared to pooled controls. (B) UMAP plot of all cells from non-regenerating and regenerating limbs colored by lineage. **(C)** Percentage of cells annotated as immune cells at each time point. **(D)** UMAP plot of subset immune cells with granular annotations. Replicate powered pseudobulk differential expression comparing limbs to regenerating time points in **(E)** T cells and **(F)** B cells. Note that all genes are statistically DE (p < 0.05) between limbs and wound healing and medium bud, but only two replicates at early bud precluded statistical comparisons. **(G)** Gene ontology of DE genes at wound healing or medium bud blastema stages as compared to limbs. GO terms filtered based on qvalue < 0.005, RichFactor > 0.5, FoldEnrichment > 5.

scRNAseq analysis during the regeneration time course uncovered an abundance of immune cells across regeneration (**Figure 1B**). Immune cell infiltration contribution peaked during wound healing, with ∼44% of captured cells being leukocytes (**Figure 1C**). Granular annotation of immune cell types using a multi-agent framework^33^ followed by manual curation uncovered a previously underappreciated diversity of innate and adaptive immune cells (**Figure 1D, Supplemental Figure 1**). While phagocytes have been implicated in limb regeneration^13^, we identified diverse and dynamic putative phagocyte clusters, including various macrophages and granulocytes such as mast cells and neutrophils (**Figure 1D, Supplemental Figure 1**). The lymphocyte compartment was more diverse than previously described^19^. We found putative natural killer (NK) cells, NK-like T cells, various T cell subsets, and both B and plasma cells (**Figure 1D, Supplemental Figure 2**), many of which have not been described in salamanders or in the regenerating limb.

We next inspected whether adaptive immune cells undergo transcriptional changes across regeneration. We found that both T and B cells across regeneration show marked transcriptional differences compared to their counterparts at homeostasis (**Figure 1E-F**). T cells showed a clear skewing away from T_H_1-like immunity, with downregulation of *Stat4*, *Ifngr1,* and *Ets1*^34^, suggesting a shift away from pro-inflammatory immune responses. Further, B cells downregulated genes important for controlling autoimmune B cell responses such as *Ets1*^35^, and anti-viral responsiveness (*Znf593*)^36^, similarly shifting away from pro-inflammatory immune responses. Consistent with these shifts in gene expression in adaptive immune cells, we observed that the myeloid compartment in the regenerating limb downregulates antigen presenting machinery, genes related to stimulating T cell mediated immunity, and cytokine and interferon responses (**Figure 1G**). Such responses are rapid, as by wound healing stage (i.e., 72 hours post amputation) these shifts in gene expression have already occurred. Altogether, our findings demonstrate that the adaptive immune system is engaged during regeneration and rapidly shifts away from inflammation towards immunosuppression.

### The blastema niche is immunosuppressed

We next sought to functionally test whether the blastema was indeed an immunosuppressive niche. To assess the immunosuppressive capacity of the regenerating limb environment, we allografted skin from transgenic donor expressing *green fluorescent protein* (GFP) under the control of a ubiquitous *CAG* promoter (*CAG:loxP-GFP-loxP-Cherry*^Simon^, hereafter referred to as GFP-positive newts)^37^ directly into the medium bud stage blastema of regenerating newt forelimbs **(Figure 2A**). To avoid potential systemic immunosuppressive effects^23^, as a control, an equivalent allograft was performed in which GFP-expressing skin was allografted onto the homeostatic hind limb of a newt regenerating both forelimbs at the medium bud stage **(Figure 2A**). Longitudinal imaging was performed to monitor graft rejection rates between the two conditions.

**Figure 2:**
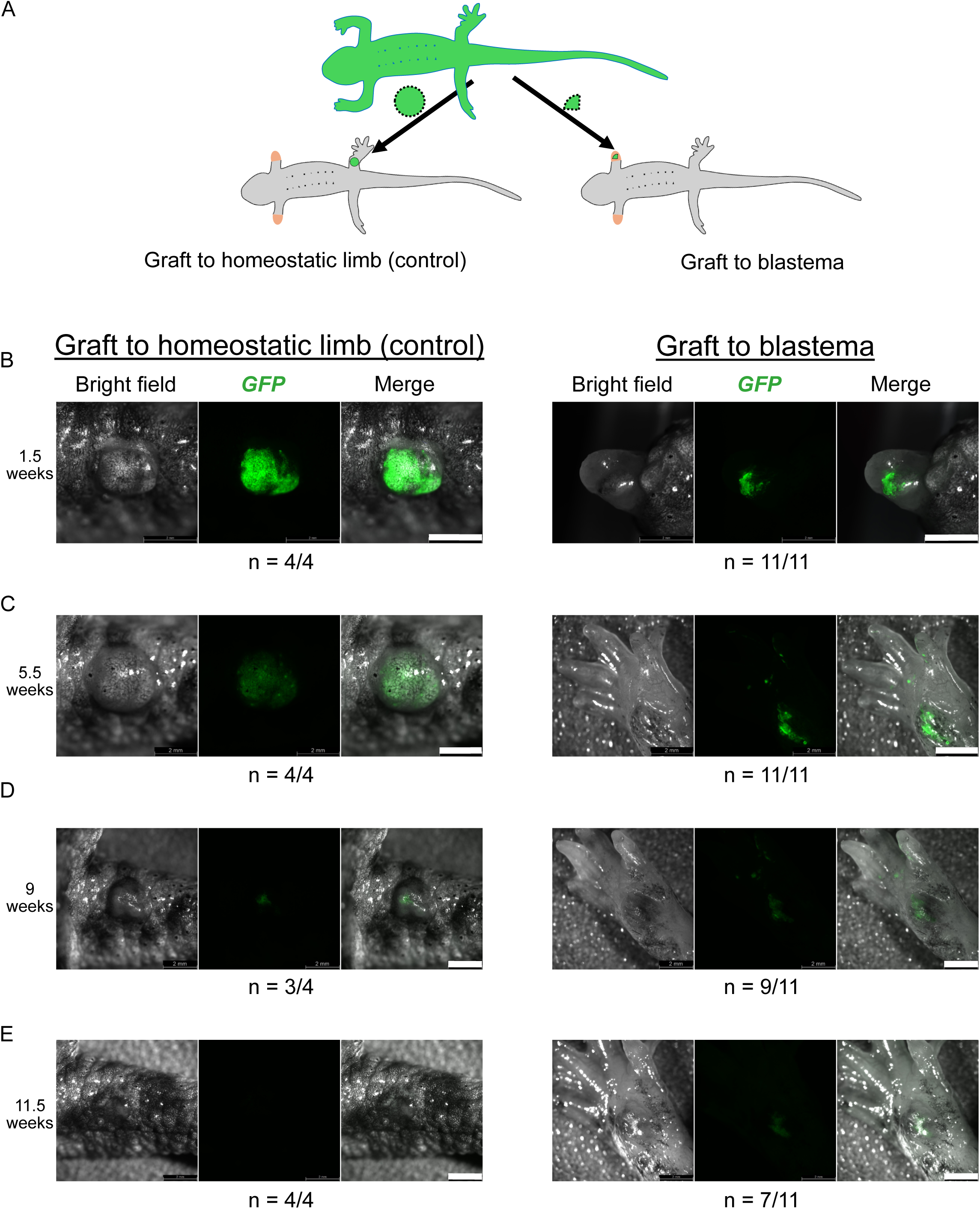
The blastema niche is immunosuppressed. **(A)** Schematic showing graft preparation from a single GFP skin donor to either homeostatic control hindlimbs or forelimb blastemas. Green circle and green quadrant signifies the sizes of GFP+ skin grafted on the intact control limb and in the blastema respectively. All animals in both control and experimental groups received bilateral forelimbs amputation and developed blastemas by the time of grafting. **(B - E)** Representative images in bright field, GFP screen and overlay (merge) of visualized graft locations on the same grafted animals at **(B)** 1.5 weeks post transplantation. **(C)** 5.5 weeks, **(D)** 9 weeks, **(E)** 11.5 weeks. “n” signifies the fraction of animals in each group that are represented by the image. Scale bars = 2mm.

GFP fluorescence in the control grafts at 5 weeks after transplantation underwent a marked reduction, indicating allograft rejection **(Figure 2C**). In contrast, GFP signal within blastema grafts was maintained over the same interval. Further, GFP fluorescence in the newly regenerated limb **(Figure 2C**) remained comparable to the initial signal post transplantation **(Figure 2B**) and was readily detectable within the regenerating limb. Additionally, the GFP-positive tissue was not merely retained at the site of transplantation but integrated progressively into the regenerating limb **(Supplemental Figure 3)**, and as expected, contributed to regeneration resulting in dispersion of graft tissue into the regenerated limb **(Figure 2C).** In contrast, control grafts onto hindlimbs were stationary.

Following completion of limb regeneration, GFP fluorescence within the blastema-derived grafts gradually declined. Nevertheless, GFP-positive tissue remained detectable in 7 of 11 animals (63%) at 11 weeks post transplantation, despite regeneration having ceased before this time point **(Figure 2D-E**). Whereas in the control animals, GFP signal was progressively rejected **(Figure 2D**) and was no longer detectable (0/4) at this same time point **(Figure 2E, Supplemental Figure 4)**.

Together, these findings indicate that the regenerative environment permits allograft persistence and survival, corroborating transcriptomic evidence that the regenerative niche is immunosuppressed. They support a model in which the blastema constitutes an immunosuppressed environment capable of accommodating allogeneic tissue during regeneration and motivate the investigation into the role of adaptive immunity in complex multi-tissue regeneration.

### Identification of the newt ortholog of *Recombination activating gene 1* (*Rag1*)

The marked transcriptional shifts in adaptive immune cells during regeneration and the immunosuppressed regeneration niche prompted us to investigate if adaptive immunity played a role during regeneration. The adaptive immune system depends on the production of a diverse repertoire of antigen receptors on the B and T lymphocytes via RAG mediated recombination^38–41^. Disruption of the *Rag1* results in immunodeficiency in jawed vertebrates, characterized by the absence of mature B and T lymphocytes^42–44^.

Given that adaptive immune responses become more robust following metamorphosis, we pursued functional experiments in the naturally metamorphosing Iberian ribbed newt (*P. waltl*). Using the recently assembled newt genome^45^, we identified and characterized for the first time the putative newt *Rag1* gene ortholog. Assessing synteny across multiple vertebrate genomes, including human, mouse, axolotl, and *Xenopus* confirmed the conserved genomic context of the *Rag1* locus (**Figure 3A**). In addition, the overall domain organization of the protein sequences was conserved across the featured species (**Figure 3B**). Notably, pairwise structure alignment^46^ of *P. waltl* Rag1 showed retained secondary structure predictions, and preservation of the core RAG1 fold (**Supplemental Figure 5A-E**). These observations indicate that the structural framework of *Rag1* gene and its protein product are conserved among vertebrates.

**Figure 3:**
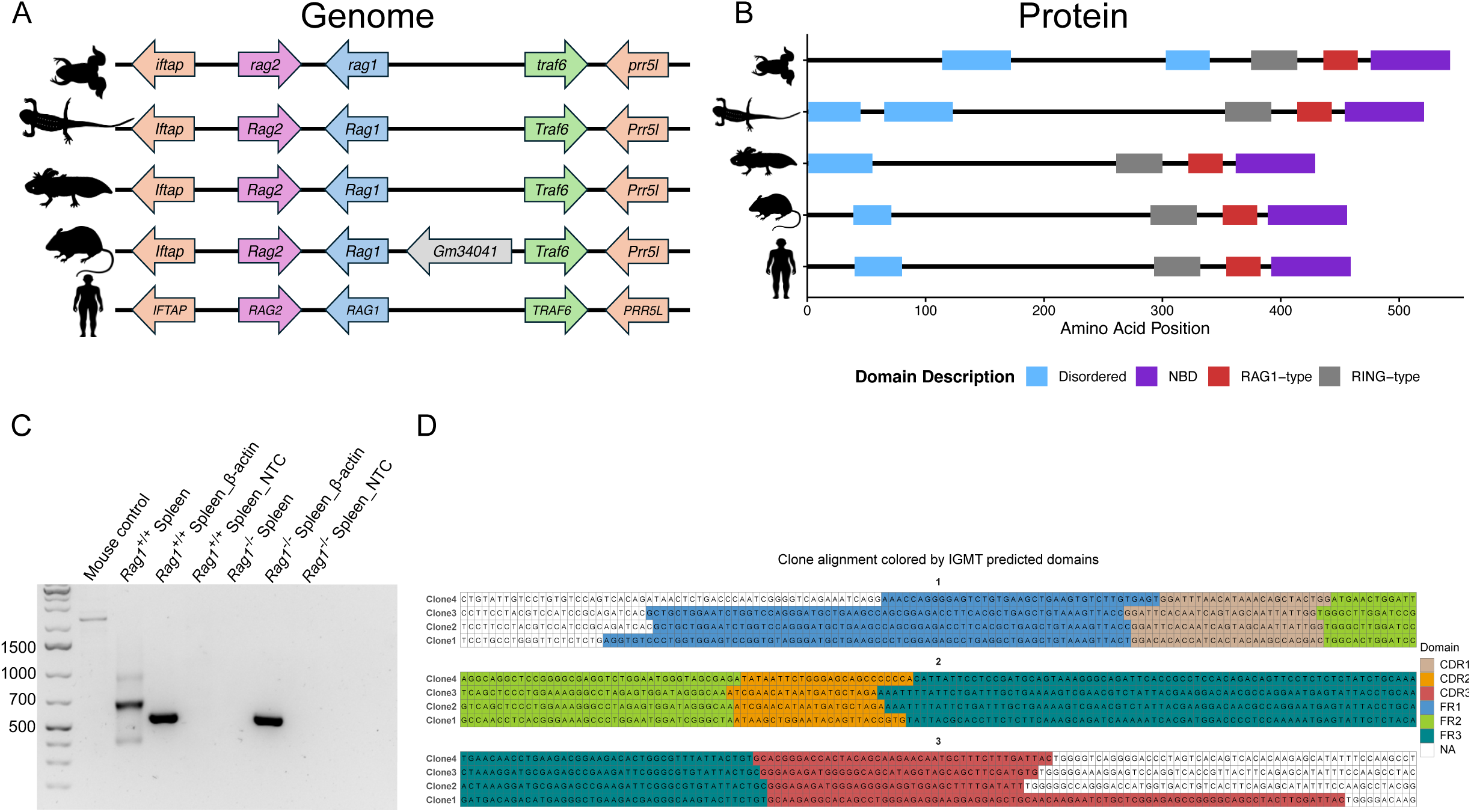
Generation and characterization of *Rag1-/-* newts. **(A)** Syntenic block illustration of gene environment across compared species; from the top, *X. tropicalis*, *P. waltl*, *A. mexicanum*, *M. musculus*, and *H*. *sapiens*. Right and left- pointing arrows represent gene location on forward and reverse strands respectively, gene length not shown in illustration. Syntenic data illustration adapted from NCBI genome data of the reported species. **(B)** Rag1 protein domain conservation across the featured species. Domains described in the legend are unanimous to the protein orthologs available on UniProtKb database; A0A8J0QTA5 for *X. tropicalis*, A0AAV7U3D9 for *P. waltl*, A7KS68 for *A. mexicanum*, P15919 for *M. musculus*, and P15918 for *H*. *sapiens*. Plots were generated using the Jsonlite R package (version 2.0.0). **(C)** 5’ RACE PCR products from splenic immunoglobulin mRNAs in *Rag1*^+/+^ and *Rag1*^-/-^ *P.waltl*. RACE reaction amplified immunoglobulin cDNAs from the constant region in the 5’ direction bearing the V[D]J components. RACE products were seen in *Rag1*^+/+^ spleen, whereas no RACE product was observed in *Rag1*^-/-^ spleen. PCR for β-actin in the template cDNA for the *Rag1*^+/+^ and *Rag1*^-/-^ spleens gave products of expected sizes as cDNA quality control. **(D)** IgBLAST and alignment of immunoglobulin cDNA sequences from four randomly selected clones of the *Rag1*^+/+^ RACE products, showing various immunoglobulin domains identifiable in the newt.

### Generation of *Rag1*-mutant *P. waltl* via genome editing

We next sought to test the role of adaptive immunity in regeneration by disrupting *Rag1* function *in vivo*. We targeted *Rag1* in one-cell-stage *P. waltl* embryos using CRISPR/Cas9-mediated genome editing. As a visible marker for successful delivery of Cas9 ribonucleoprotein complexes (RNPs), we also targeted the *Tyrosinase* gene^47^, which encodes a key enzyme in melanin biosynthesis. Loss of pigmentation seven days post-injection provided rapid confirmation of RNP activity (**Supplemental Figure 6D**). All embryos injected with *Tyrosinase*-targeted sgRNAs exhibited complete depigmentation (100%, n = 29), validating efficient CRISPR/Cas9 delivery.

### *Rag1* deficiency abolishes V(D)J recombination in *P. waltl*

We analyzed editing efficiency in the embryos to confirm targeted disruption. These first-generation (F_0_) mutants exhibited varying degrees of editing at the *Rag1* locus, with heterogeneous indel profiles and predicted knockout efficiencies (**Supplemental Figure 7A**). Given this variability, these F_0_ animals cannot be uniformly classified as *Rag1*^-/-^ and are referred to as “crispants” throughout this study. Using these founders, we bred two stable *Rag1*^-/-^ lines (**Supplemental Figure 7B**), one with a -1 base pair (bp) deletion at position 1005, and another with a -7bp deletion at position 1004 of the transcript (**Supplemental Figure 7C**). These two putative knockout lines now allow for the characterization of *Rag1* deficiency on immune function and regeneration.

We sought to validate the functional loss of RAG1-mediated recombination activity to confirm functional gene knockout. The RAG1 protein is essential for initiating somatic recombination of the variable (V), diversity (D), and joining (J) gene segments at immunoglobulin and T cell receptor (TCR) loci, enabling the generation of a diverse repertoire of B and T cell receptors. In the absence of functional RAG1, V(D)J recombination fails, resulting in an absence of mature B and T lymphocytes due to non-rearranged antigen receptor loci^48^. To assess V(D)J recombination activity, we performed 5′ rapid amplification of cDNA ends (5′ RACE-PCR) on immunoglobulin mRNAs isolated from the spleen of WT and *Rag1*^-/-^ newts^49^. This approach targets the 5′ region upstream of the constant region, where the V(D)J segments are joined during lymphocyte development. RACE-PCR products were readily detected by gel electrophoresis in WT samples, indicating successful recombination and expression of rearranged antigen receptor transcripts (**Figure 3C**). In contrast, no RACE-PCR products were observed in *Rag1^-/-^* spleen, consistent with an absence of V(D)J recombination (**Figure 3C**). Amplification of β-actin from the same cDNA confirmed that the lack of RACE products in *Rag1*^-/-^ reflected absence of recombination rather than poor sample quality (**Figure 3C, Supplemental Figure 8A**).

To further characterize the recombination events, we cloned and sequenced the RACE-PCR products from WT spleen. IgBLAST^50,51^ run on all the selected clones confirmed domains consistent with established immunoglobulin sequences in the complementarity-determining region 3 (CDR3) among individual clones (**Figure 3D**) and alignment of the clone sequences presented distinct recombination events among individual clones (**Supplemental Figure 8B**). These results provide molecular evidence of V(D)J recombination in WT animals and its absence in *Rag1*^-/-^ newts, confirming the functional inactivation of adaptive immune recombination machinery, and particularly, a deficiency in immunoglobulin producing B cells.

### *Rag1^-/-^* newts have markedly reduced T cells

The lack of antigen receptor recombination should result in the absence of mature B and T cells. Given *Rag1*^-/-^ animals did not demonstrate recombination of their B cell receptor (**Figure 3C**), we chose to investigate T cell numbers to test whether lack of V(D)J recombination resulted in decreased adaptive immune cells. We assessed T cell presence in spleen sections by immunofluorescent staining for the pan T cell marker, CD3. First, we obtained that spleen sizes in *Rag1*^−/−^ newts were strikingly smaller than spleens from WT and *Rag1*^+/−^ newts of comparable sizes (Cohen’s *d* = 2.39) (**Figure 4A-B**). In WT and *Rag1*^+/−^ controls, CD3⁺ cells were abundant, and distributed throughout the spleen (**Figure 4C-D**). In contrast, CD3^+^ cells were markedly reduced in *Rag1*^−/−^ newt spleens, with signal levels comparable to negative controls (**Figure 4C-D**). Quantification of cell count per section revealed a marked reduction in CD3^+^ cells in spleens from *Rag1*^−/−^ newts (Cohen’s *d* = 2.56) (**Figure 4E**). These findings are consistent with other *Rag1*^−/−^ models^42,52^ and verify the loss of T cells in *Rag1*^−/−^ newts.

**Figure 4:**
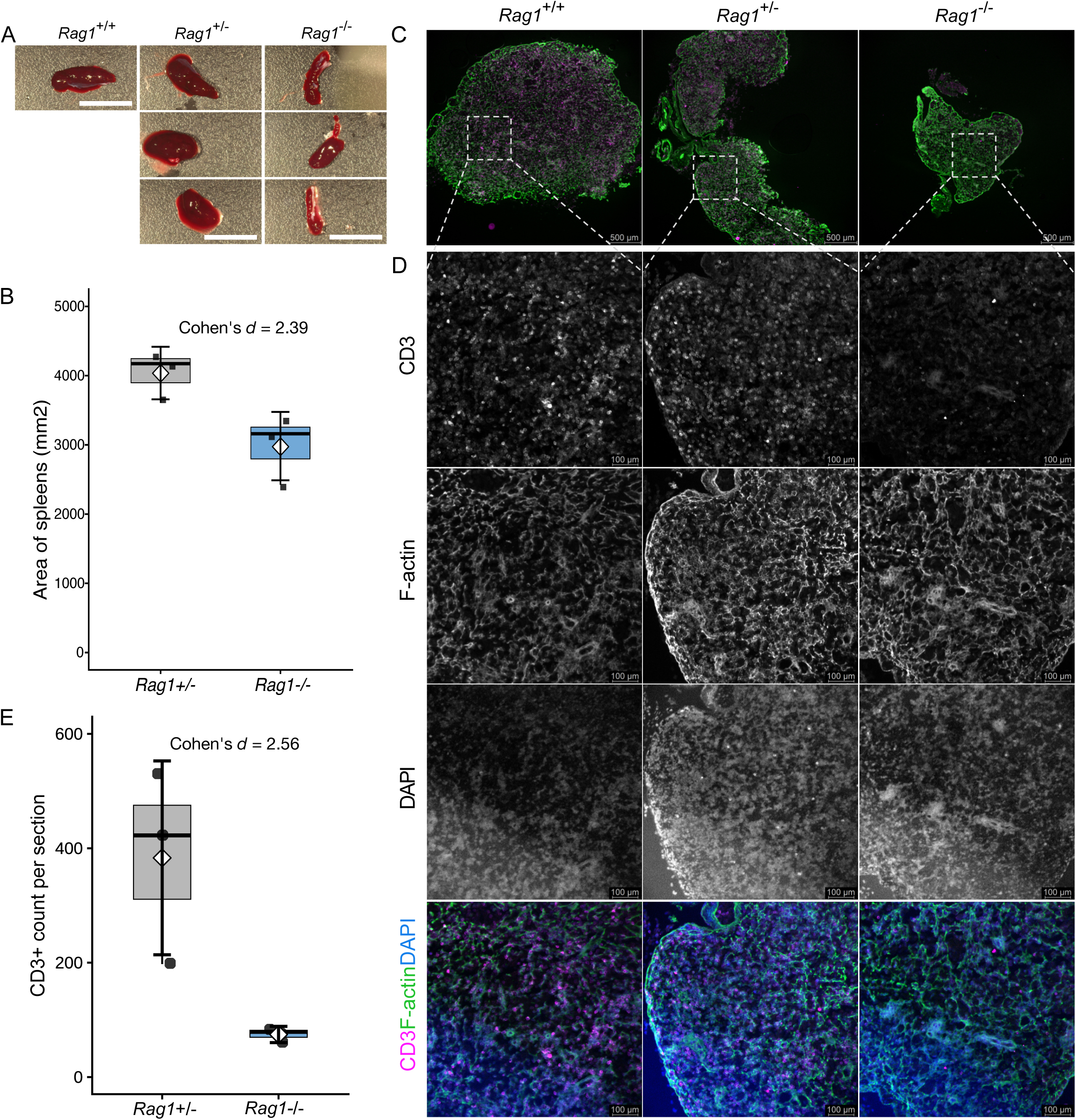
*Rag1^-/-^* newts have T cell deficiency. **(A)** Spleen images from size-matched WT, *Rag1*^+/−^, and *Rag1*^−/−^ newts visualizing size differences (scale bar, 10 mm). **(B)** Visual quantification of spleen size differences between *Rag1*^+/−^ and *Rag1*^−/−^ newts. Quantification measured by pixel area covered by the spleen obtained from ImageJ; Cohen’s *d* = 2.39. **(C-D)** Representative immunofluorescence images of spleen sections (12 μm) from post-metamorphic *P. waltl* animals. Sections were stained for CD3 (Alexa Fluor 647, cyan) to label T cells and with phalloidin–Alexa Fluor 488 (green) to visualize F-actin; nuclei were counterstained with DAPI (blue). Individual fluorescence channels are shown in grayscale to highlight signal intensity. Merged images are displayed with corresponding false colors. **(C)** Low-magnification overview of the spleen (scale bar, 500 μm). **(D)** Higher-magnification view of the indicated region (scale bar, 100 μm). Images are representative of WT (n=1), *Rag1*^+/−^ (n = 3), and *Rag1*^−/−^ (n = 3). **(E)** Quantification of average CD3+ cells per immunofluorescent labeled spleen section from *Rag1*^+/−^, and *Rag1*^−/−^ newts; Cohen’s *d* = 2.56. Cohen’s *d* effect sizes were interpreted as small (∼0.2), medium (∼0.5), and large (≥0.8).

### *Rag1*^-/-^ leads to impaired survival

Since *Rag*-associated immunodeficiency leads to increased susceptibility to infections and other immune dysregulations, we monitored the long-term survival of both F_0_ crispants and F_2_ knockout lines. To evaluate the potential impact of *Rag1* disruption on survival in crispants, we calculated knockout scores for each animal and locus, and binned crispants into two groups: *Rag1_*lowKO (knockout score 0–89) and *Rag1_*highKO (≥90 KO score at either sgRNA site or a cumulative score ≥140 across both sites) (**Supplemental Figure 7A**). Median overall survival was significantly shorter in the *Rag1_*highKO group (376 days; 95% CI: 230–460) compared to the *Rag1_*lowKO group (460 days; 95% CI: 460–474, *p*-values=5 × 10^−4^) and dechorionated controls (460 days; 95% CI: 460–460, *p*-values <1 × 10^−4^) (**Figure 5A**). Our findings suggest that high-efficiency *Rag1* disruption is associated with significantly impaired survival, consistent with the predicted immunodeficient phenotype^52^.

**Figure 5:**
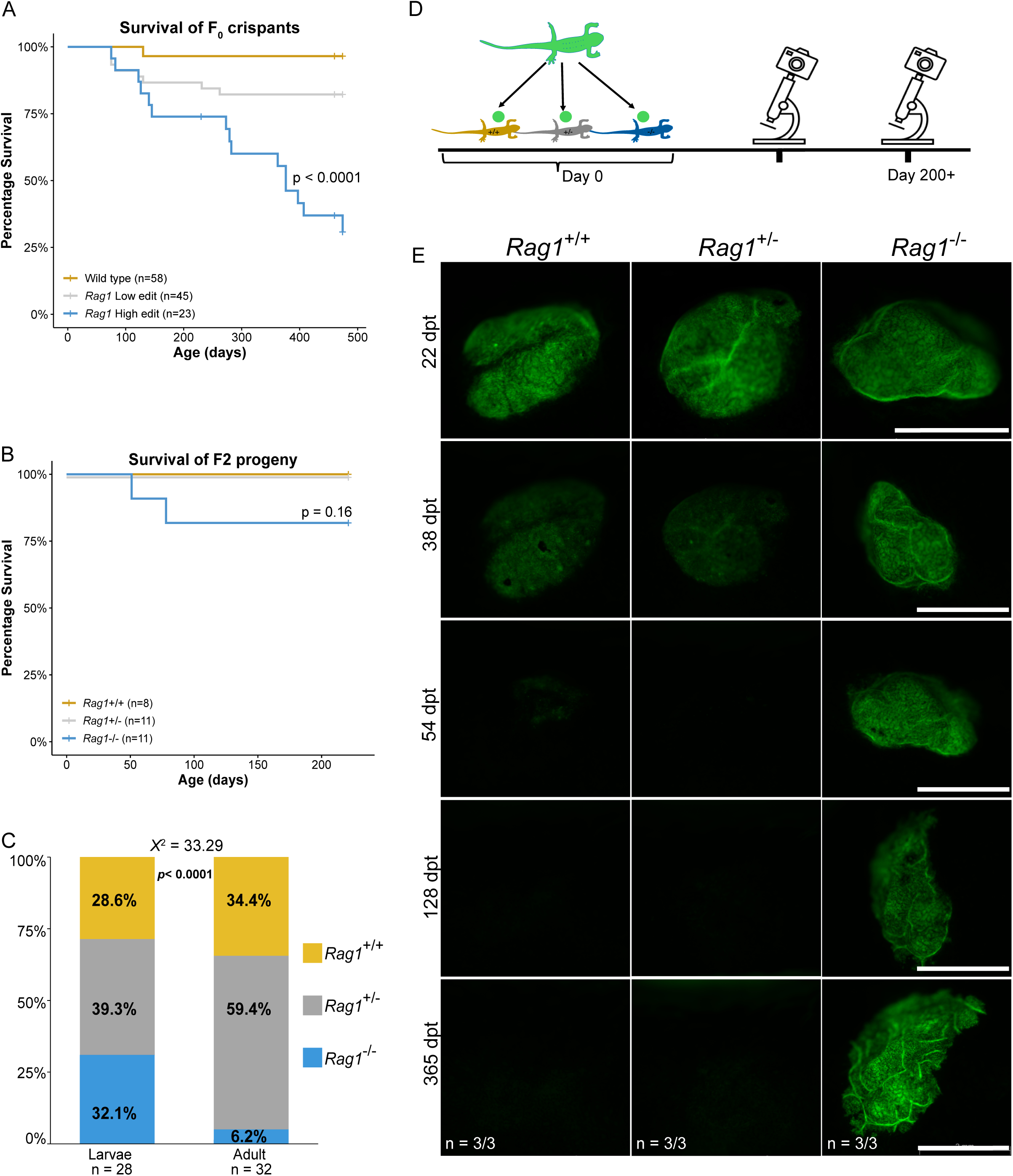
*Rag1^-/-^* newts are immunodeficient. **(A)** Kaplan-Meier curve of the survival of “*Rag1* High edit” (blue) in comparison to “*Rag1* Low edit” (grey) and “WT” (yellow) newts. A log-rank test showed a significant difference between “WT” and “*Rag1* High edit” at 95% C.I. of 7.5 - 80.3 (*p* < 0.0001) and hazard ratio 24.7. **(B)** Kaplan-Meier curve of the survival of F_2_ *Rag1*^-/-^ (blue) in comparison to *Rag1*^+/-^ (grey) and *Rag1*^+/+^ (yellow). **(C)** Stacked bar plot of genotypes from offspring from crosses of *Rag1*^+/-^ parents as larvae (left) or adults (right). **(D)** Transplantation procedure showing grafts prepared from a single donor and grafted onto *Rag1*^+/+^, *Rag1*^+/-^ and *Rag1*^-/-^ animals, which remained under extensive continuous monitoring. **(E)** Representative images of graft locations on the same grafted animals. Green fluorescence indicates graft persistence on the animal at the recorded time points; shown here are 22 dpt, 38 dpt, 54 dpt, 128 dpt; and 365 dpt n = 3/group in repeated experiments, scale bar = 2mm.

To confirm this finding in F_2_ *Rag1*^-/-^ mutant newts, we followed a cohort of F_2_ *Rag1*^-/-^ newts over ∼200 days and, like the highly edited F_0_ crispants, we observed some lethality around metamorphosis (∼75 days post-fertilization) but no significant differences in survival of *Rag1*^-/-^ compared to WT siblings (**Figure 5B**). While F_2_ survival study was relatively shorter than the F_0_ survival study, this difference reflects the period F_0_ animals were raised to reach sexual maturity to produce F_1_ offspring and further generations. We next evaluated the impact of *Rag1* disruption on survival by assessing Mendelian ratios of F_2_ larval and adult newts. We genotyped 28 randomly selected larval F_2_ individuals, which revealed predicted Mendelian inheritance of the *Rag1* allele: 8 animals (28.6%) were WT, 11 (39.3%) were heterozygous, and 9 (32.1%) were homozygous for the mutant allele (**Figure 5C**). These results show that *Rag1*^-/-^ salamanders developed normally through embryogenesis and early development. However, genotyping a cohort of 32 randomly selected adult F_2_ animals that were not previously genotyped as larvae revealed a selection against the *Rag1* allele: 11 animals (34.4%) were WT, 19 (59.4%) were heterozygous, and 2 (6.2%) were homozygous for the mutant allele, with this distribution differing significantly from the larval cohort, *X*^2^ = 33.29, *p* < 0.0001 (**Figure 5C**). The deviation from expected Mendelian inheritance in adults suggests that *Rag1*^-/-^ newts post metamorphosis have a survival disadvantage compared with WT and heterozygous newts. Paired with the *Rag1_*highKO crispants worse survival, our results show that *Rag1* knockout impairs survival likely due to immunodeficiency.

### *Rag1^-/-^* newts are immunodeficient as evidenced by the inability to reject allografts

To functionally validate the immunodeficiency of *Rag1*^-/-^ newts, we assessed their ability to reject allogeneic skin grafts. To indelibly mark allografted skin, we used a transgenic donor expressing GFP. Circular full-thickness skin punch biopsies (3 mm in diameter) were excised from the tail of the GFP-positive newts and grafted onto *Rag1*^+/+^, *Rag1*^+/-^, and *Rag1*^-/-^ allogeneic recipient newts. GFP-positive grafts were imaged serially after transplantation to monitor the onset and progression of immune-mediated rejection (**Figure 5D**). Complete graft rejection, defined by the clearance of donor GFP-expressing skin cells, occurred in chronic fashion in WT and heterozygous newts between 38 and 52 days post-transplantation (dpt) (**Figure 5E**). In contrast, *Rag1*^−/–^ animals retained GFP-positive donor skin cells without detectable rejection, persisting throughout the study and up to 365 dpt (**Figure 5E**). These results demonstrate the chronic allograft rejection response characteristic of newts is adaptive immune mediated. Further, the inability to reject allogeneic grafts demonstrates that *Rag1*^−/–^ newts are functionally immunodeficient.

### Pre-metamorphic *P. waltl* appendage regeneration occurs independently of adaptive immunity

Having established that *Rag1*^-/-^ salamanders lack functional adaptive immunity, we next investigated whether adaptive immunity is required for appendage regeneration in pre-metamorphic *P. waltl*. Age-matched, sibling, pre-metamorphic *Rag1*^-/-^ and WT animals were subjected to limb or tail amputation and monitored throughout regeneration. Both groups formed a morphologically indistinguishable blastema, and outgrowth progressed with comparable kinetics (**Figure 6A-B**). By 24 days post amputation (dpa), regenerated limbs in *Rag1*^-/-^ animals were indistinguishable from their WT counterparts in terms of morphology and size (**Figure 6A**). These data indicate that adaptive immune cells are dispensable for successful appendage regeneration in pre-metamorphic newts.

**Figure 6:**
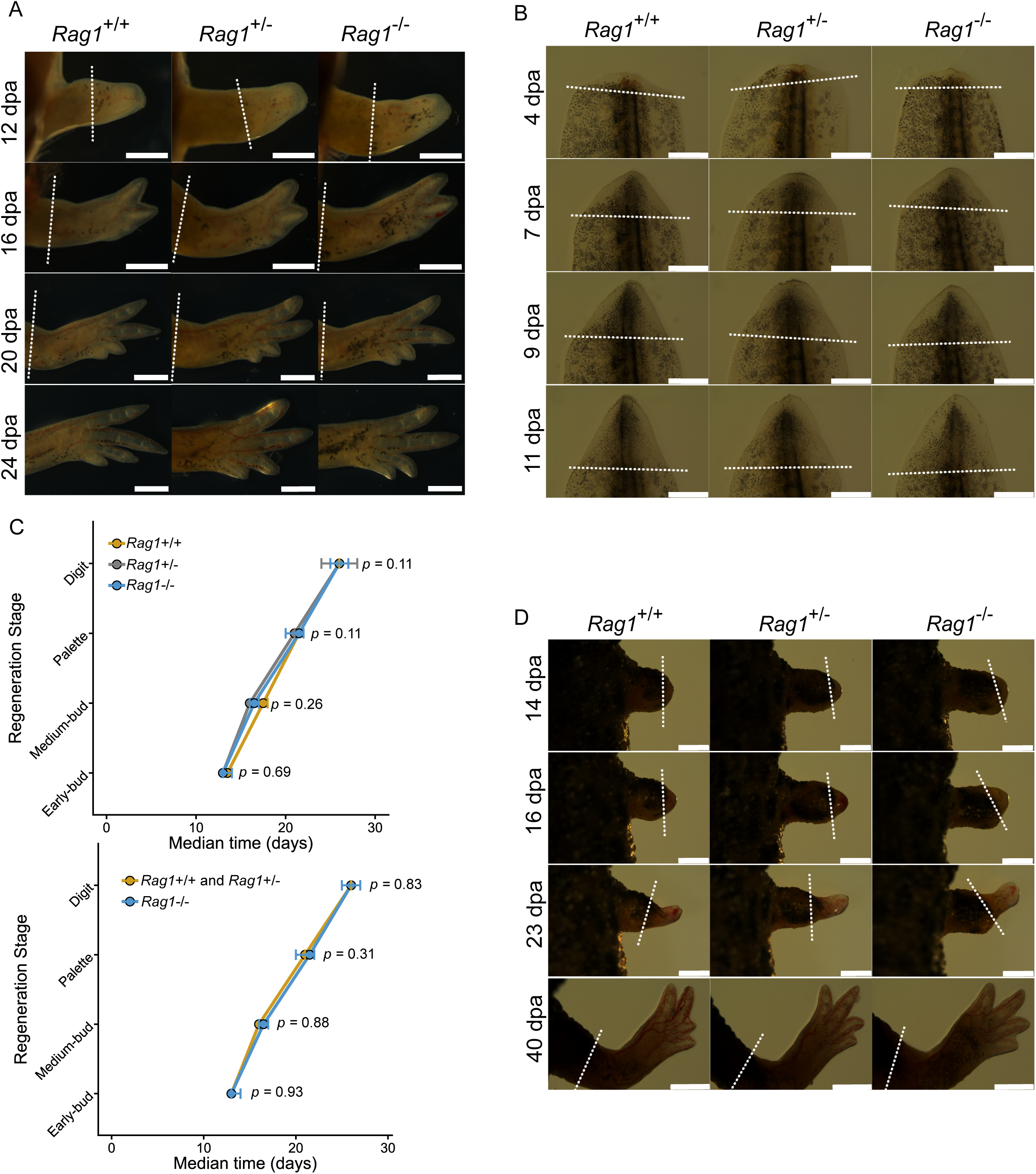
Adaptive immunity is dispensable for appendage regeneration in pre- and post-metamorphic *P. waltl*. **(A)** Representative images of pre-metamorphic limb regeneration timeline across *Rag1*^+/+^ (n = 2), *Rag1*^+/-^ (n = 9) and *Rag1*^-/-^ (n = 5) animals from the palette stage to the full limb regenerates. The regenerates showed no favored or impeded outcomes across the genotype groups. The white dashed lines indicate plane of amputation; scale bars = 2mm. **(B)** Representative images of pre-metamorphic tail regeneration timeline across *Rag1*^+/+^ (n = 5), *Rag1*^+/-^ (n = 17) and *Rag1*^-/-^ (n = 2) animals from the blastema stage to > 80% tail restoration. Similar regenerate outcomes across the genotype groups. The white dashed lines indicate plane of amputation; scale bar = 1mm. **(C)** Post-metamorphic regeneration shows no significant difference in the median times to reach regeneration events in two different scenarios. Upper panel: *p* = 0.69 at early-bud blastema, *p* = 0.26 at medium-bud blastema, *p* = 0.11 at palette, *p* = 0.11 at digit differentiation, *Rag1*^+/+^ (n = 5), *Rag1*^+/-^ (n = 6) and *Rag1*^-/-^ (n = 8). Lower panel: *p* = 0.93 at early-bud blastema, *p* = 0.88 at medium-bud blastema, *p* = 0.31 at palette, *p* = 0.83 at digit differentiation. n = 11 and 4 respectively for *Rag1*^+/+^+*Rag1*^+/-^ and *Rag1*^-/-^. **(D)** Representative images of post-metamorphic limb regeneration timeline across *Rag1*^+/+^ (n = 2), *Rag1*^+/-^ (n = 9) and *Rag1*^-/-^ (n = 4) animals from the blastema stage to full limb regenerates. Similar regenerate outcomes across the genotype groups. The white dashed lines indicate plane of amputation; scale bars = 2mm. This experiment was repeated twice with similar outcomes, and n numbers here reflect one individual experiment.

### Adaptive immunity is dispensable for adult newt limb regeneration

Given that amphibian metamorphosis is accompanied by extensive physiological remodeling, including reorganization of the immune system^53^, we next assessed regeneration capacity of adult newts. We amputated sibling *Rag1*^−/–^ and WT animals that had undergone metamorphosis. At 19 dpa, both genotypes exhibited comparable blastema formation (**Figure 6C**). Regeneration proceeded to completion without overt morphological differences in regenerated limbs, regardless of genotypes (**Figure 6D**). These results demonstrate that throughout life adaptive immunity is dispensable for appendage regeneration in newts.

## Discussion

With the growing interest at the interface of regeneration and immunology, the role of adaptive immunity in appendage regeneration has remained enigmatic. Our study, and that of Bolanos Castros *et al*.^54^, provide definitive evidence that adaptive immunity is dispensable for appendage regeneration in larval and adult salamanders. This substantially advances our understanding of the required immune inputs to appendage regeneration. Further, this introduces a unique immunodeficient salamander line that provides a novel platform to explore previously unattainable experiments in salamanders (e.g., xenotransplantation).

There is a dearth of studies on adaptive immunity in complex tissue regeneration. Previous work has demonstrated a requirement for Tregs in regeneration of spinal cord, heart, and retina in zebrafish^55^ and the mouse digit tip^56^. Our findings in *Rag1*^-/-^ animals suggest that Tregs are not required for appendage regeneration in newts. However, it does not rule out the possibility that Tregs are required in the presence of the full arsenal of adaptive lymphocytes. It is important to note that lymphoid deficient (i.e., NOD scid gamma - NSG) mice regenerate normally, implying that lymphoid cells are dispensable in mouse digit tip regeneration^57^. Altogether, this presents a complicated role for lymphocytes in regeneration, as the balance of cytotoxicity and immunosuppression appear to be critical in dictating regenerative outcomes. One potential reason why cytotoxic lymphocytes would interact with regenerative tissue would be as a means of quality control, similar to the role of macrophages in hematopoiesis^58^. Thus, Treg depletion would result in overexuberant cytotoxicity to progenitors and thus impair regeneration—a phenotype that would only manifest in immune replete animals. Further studies using conditional depletion strategies (e.g., NTR2.0^59^) will be instrumental in teasing apart individual immune cells’ role in appendage regeneration.

The noted anti-correlation between mammalian-like adaptive immune responses and regenerative capacity suggests that adaptive immunity may impede regeneration^60^. Salamanders are known for their chronic graft rejection responses^61^. Whether this limited immunological function reflects an evolutionary balance in which reduced adaptive immune capacity ensures regenerative success is yet to be determined. A better understanding of the origins of newt’s adaptive immune chronicity could provide a means to enhance newt’s adaptive immune responses post-amputation and test compatibility with regeneration.

The dispensable nature of adaptive immunity positions *Rag1*^-/-^ newts as a major asset to study regeneration. While many studies have relied on allogeneic transplant to learn about lineage relationships^7,62^, it may be worth revisiting these experiments in the absence of potential confounding allogeneic anti-graft responses. In addition, the inability to reject foreign material will allow for previously impossible xenotransplantation experiments. Revisiting previously failed attempts of transplanting cells from species that lose regeneration capacity with age (e.g., *Xenopus*)^63^ will provide unique insights into mechanisms of regeneration and lack thereof.

While our data establish that *Rag1*-dependent adaptive immunity is not required for salamander limb regeneration, this study does not exclude compensatory roles for innate or innate-like lymphoid populations that may substitute for adaptive immune functions. Our analyses focused on gross regenerative outcomes and may not capture more subtle defects in patterning, cellular composition, or tissue function. Given the important role T cells have been shown to play in heart regeneration^26,55^, it will be important to explore other regenerative paradigms in *Rag1* deficient newts.

An abundance of evidence in mammals suggests that adaptive immune responses can aggravate injuries and contribute to poor regenerative outcomes^64^. Our transcriptomic immunophenotypes and blastema grafting experiment add a new layer to our understanding of blastema biology—that the blastema represents an immunosuppressive milieu. This leaves open the major question of whether immune suppression is necessary to control anti-regenerative adaptive immune responses. It will be critical to elucidate these mechanisms of immune restraint to understand how immune responses diverge between regenerative and non-regenerative species and determine what constitutes a pro-regenerative immune milieu. Harnessing this knowledge may guide the rational development of regenerative therapies by transiently attenuating adaptive immune activity or engineering immune environments that emulate the permissive conditions characteristic of highly regenerative vertebrates.

## Materials and Methods

### Animal husbandry: breeding and maintenance

All procedures in this report regarding animal handling, care and treatment were performed in accordance with the guidelines approved by the Swedish regulations (Jordbruksverket) under the ethical permit number *5.8.18-16345/2021*. The *P. waltl* used in this study were from the colony at Lund University. The animals were reared at 12 hours light/12 hours darkness at 19–22°C in carbon filtered tap water augmented with 55g Tetra marine salt, 15g of Ektozon salt, and 2.5mL of Prime^®^ water dechlorinator per 100L of water. Animals were narcotized where applicable in 0.1% of Tricaine (MS222 Sigma) in conditioned tap water (pH 7.0). Once completely narcotized, animals were promptly taken through the proposed procedure. After the procedure, when not terminal, animals were then transferred back to the container with conditioned tap water (pH 7.0); all non-terminal animals were allowed to recover and regenerate according to experimental plans to assess for regeneration phenotypes.

### Single cell RNA sequencing

Data were reprocessed from Leigh *et al*. 2018^19^. All code for reprocessing are available on Github: https://github.com/RegenImm-Lab/adaptive_immune_limb_regen. Data were aligned to the axolotl genome (GCF_040938575.1) using a previously modified inDrops pipeline (https://github.com/brianjohnhaas/indrops). We summed isoforms to the gene level and ran output matrices through Cellbender^65^. Seurat^66^ was used to analyze data and create UMAPs as per the deposited code. In brief, we created per sample count matrices, filtered cells (nFeature_RNA > 200 and < 3750–4500 percent.mt < 5–8%, thresholds varying per batch) and merged batches using Canonical Correlation Analysis (CCA). We used default FindAllMarkers to identify cluster specific markers to feed into CyteTypeR^33^ for cell annotation. This was enhanced by providing gene description information when only locus IDs were available in the gff file. We then subset immune cells and re-annotated cells using the same steps as for the full dataset (i.e., FindAllMarkers, provide gene descriptions into CyteTypeR). Using these cell annotations, we calculated immune cell abundance at the different regenerative stages. We performed biological replicate powered pseudobulk differential expression analysis using edgeR^67^. Differentially expressed genes were carried forward into Gene Ontology (GO) analysis, and we displayed filtered (qvalue < 0.005, RichFactor > 0.5, FoldEnrichment > 5) results for abundant cell types: T cells, Phagocytic Macrophages, Macrophage, Neutrophil−Like Granulocyte. To aid Gene Ontology analysis we performed functional annotation of axolotl proteome using EggNog like so: emapper.py --cpu 48 -m mmseqs --itype proteins -i GCF_040938575.1_UKY_AmexF1_1_protein.faa -o axolotl_eggnog. Since the proteome contained different isoforms, we collapsed EggNog annotations to the gene level. Using this functional annotation, we performed GO analysis using ClusterProfiler^68^. Single cell data are browsable at SciLifeLab serve: <u>axolotl-limb-</u> <u>regen-scrnaseq.serve.scilifelab.se</u>.

### CRISPR design and constitution

The *P. waltl* transcriptome^69^ and genome^45^ were used to design CRISPR-sgRNAs according to FlashFry^70^. The sgRNAs were produced by IDT. crRNA were annealed with tracrRNA and combined with Cas9 enzyme to make the ribonucleoproteins (RNPs) according to the RNP constitution protocol by Kroll *et al*^71^. 10-20nL of the constituted RNPs was injected into single cell salamander embryos.

### Microinjection

Single-cell stage salamander embryos were collected from natural spawn or induced ovulation using human chorionic gonadotropin (hCG) hormone. Embryos were de- jellied and dechorionated in 1X Marc’s Modified Ringers (MMR; 0.1 M NaCl, 2 mM KCl, 1 mM MgSO_4_, 2 mM CaCl_2_, 5 mM HEPES pH 8, 0.1 mM EDTA) water with pen-strep. Embryos were transferred to fresh 1X MMR water (at or just below room temperature) until microinjection. Microinjection for single-cell embryos were done within 4 hours of embryo collection. De-jellied and dechorionated embryos were washed in injection medium (0.75% methylcellulose preparation^72^ or 6% Ficoll solution) and transferred to fresh injection medium where they are immobilized. 10-20 nL of constituted RNPs were injected into the embryos using a glass capillary needle connected to microinjector. The embryos were allowed to stay overnight in the injection medium after microinjection and transferred to 0.1X MMR water with pen-strep at 130 units/ml. The embryos were allowed to develop and sorted from the non-developing embryos.

### Rag1 targeting

To target the functional domain of *Rag1*, we designed two single-guide RNAs (sgRNAs) (**Table 1**) spaced approximately 1,400 bp apart, with the goal of excising a critical coding region via dual-site CRISPR–Cas9-mediated cleavage (**Supplemental Figure 6A**). Both sgRNAs were co-injected as ribonucleoprotein complexes into one-cell-stage *P. waltl* embryos. Genomic DNA was subsequently extracted from tail tip biopsies, and the targeted locus was genotyped in all injected animals.

**Table 1:**
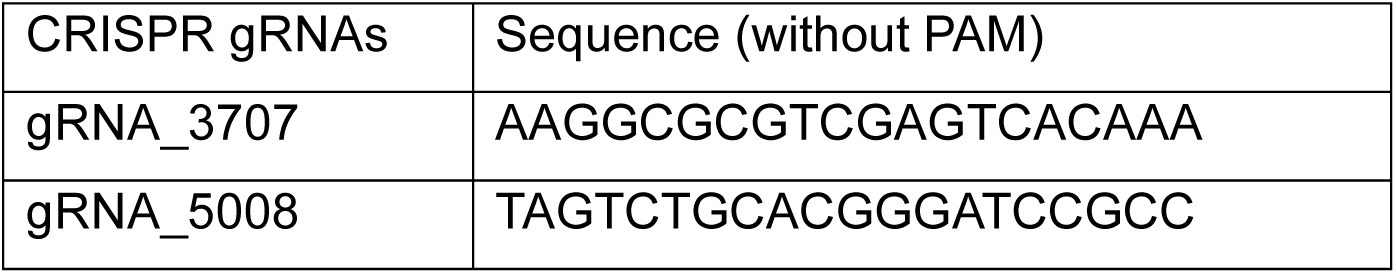
CRISPR gRNAs.

Although large-fragment excision between the two sgRNA sites was not detected, sequence analysis revealed efficient editing at the individual cleavage sites. To compare sgRNA performance, we quantified editing efficiency at each site. One sgRNA consistently induced a higher frequency of insertions and deletions (indels), with a greater proportion resulting in frameshift mutations predicted to disrupt *Rag1* open reading frame integrity (**Supplemental Figure 6B-C**). These data informed the selection of a single, highly efficient sgRNA for the generation of *Rag1*^-/-^ lines, as well as the specific mutant *Rag1* alleles retained for further analysis in this study.

### Generation of *Rag1*^-/-^ salamanders

To achieve this, we selected crispant founders that harbored frameshift-inducing mutations exclusively at the more efficient sgRNA target site, with no detectable edits at the less efficient site. These founders were outcrossed to wild-type animals from unrelated lineages. Genotyping of the F_1_ progeny confirmed successful germline transmission of the mutant *Rag1* allele. Two different allelic variants of frameshift mutations were passed to the offspring at the F_1_ progeny, viz *Rag1*^del1^ and *Rag1*^del7^, corresponding to 1bp and 7bp deletions respectively (**Supplemental Figure 7C**). We carefully curated the animals with the different variants, and only inter-crossed heterozygous carriers of the same frameshift mutation - *Rag1*^+/del1^ or *Rag1*^+/del7^ - to generate F_2_ offspring in the allelic fashion of *Rag1* ^del1/del1^ and *Rag1*^del7/del7^ as *Rag1*^-/-^salamanders (**Supplemental Figure 7B**). There were no physical or biological differences associated with the mutant allelic variations.

### Genotyping

The genomic target regions were amplified by PCR using NEB Q5^®^ High-Fidelity PCR Kit according to the kit protocol. The primers for the PCR amplification were designed around the genomic target site for a PCR product of approximately 500bp. The primers used in the PCR reactions and annealing temperatures are listed in Table 2. The PCR products were electrophoresed in 1.5% agarose gel in 1X Tris-acetate-EDTA (TAE) buffer, at 400mA for 125V for 60 minutes. The bands corresponding to 500bp were excised from the gel and cleaned. The cleaned products were sent for Sanger sequencing and the resulting ab1 files were analyzed using ICE-Synthego^73^ or TIDE^74^ (https://tide.nki.nl/) for knockout efficiencies or compared with known sequences via alignment.

**Table 2:**
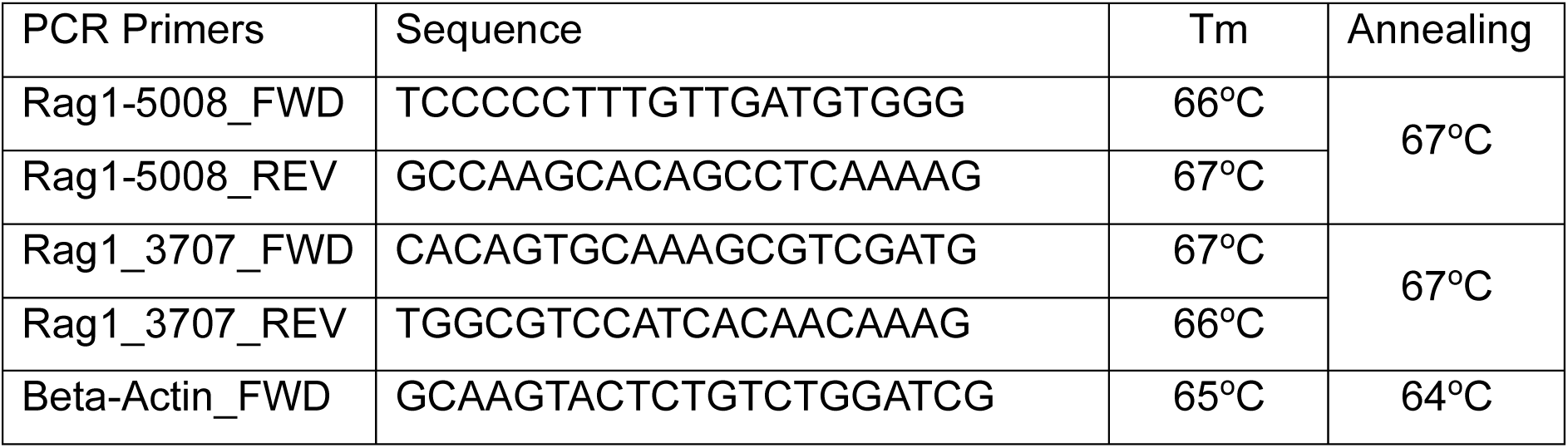
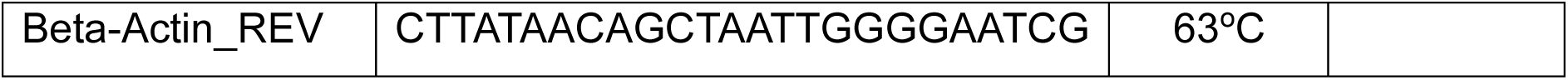
Genotyping primers.

### Amputation

Pre-metamorphic salamanders (3.5 – 5cm) were narcotized by immersion in 0.05% Tricaine and post-metamorphic salamanders (8 – 10cm) were narcotized by immersion in 0.1% Tricaine. Once completely narcotized, animal is placed in a prone position on a cast 1-2% agarose gel surface moistened with narcotizing solution. Limbs were amputated proximally at the midpoint of the humerus, and the protruding bone is trimmed off. Tails were amputated at 20% of the whole-body length (snout to tail-tip) from the tail tip. Pre-metamorphic salamanders were transferred immediately after amputation to conditioned salamander water at room temperature (20 - 22°C). Post-metamorphic salamanders were allowed to stop bleeding before transferring to conditioned salamander water.

### RACE PCR and Cloning

Total RNA was extracted individually from spleens and thymi of selected animals using the TRIzol™ Reagent, and according to manufacturer’s instructions (Pub. No. MAN0001271). Using the SMARTer RACE 5’3’ Kit (Takara Bio Inc., Japan, Cat. No. 634858) and following the manufacturer’s instructions, first strand cDNA was synthesized from the total RNA. The RACE-ready cDNA was used for PCR amplification of the 5’ ends of the cDNA using selected gene specific primers (GSPs) (**Table 3**) in accordance with the manufacturer’s instructions. The PCR products were electrophoresed in 1.5% agarose gel in 1X Tris-acetate-EDTA (TAE) buffer, at 400mA for 125V for 60 minutes. The bands were excised from the gel and cleaned and further cloned into competent Stellar cells (Clontech). Clones were picked and plasmid DNA were extracted from the clones after 16 hours of culture in LB broth medium at 37°C. The plasmid DNA were sent for Sanger sequencing, and the sequencing data were analyzed on Snapgene Viewer.

**Table 3:**
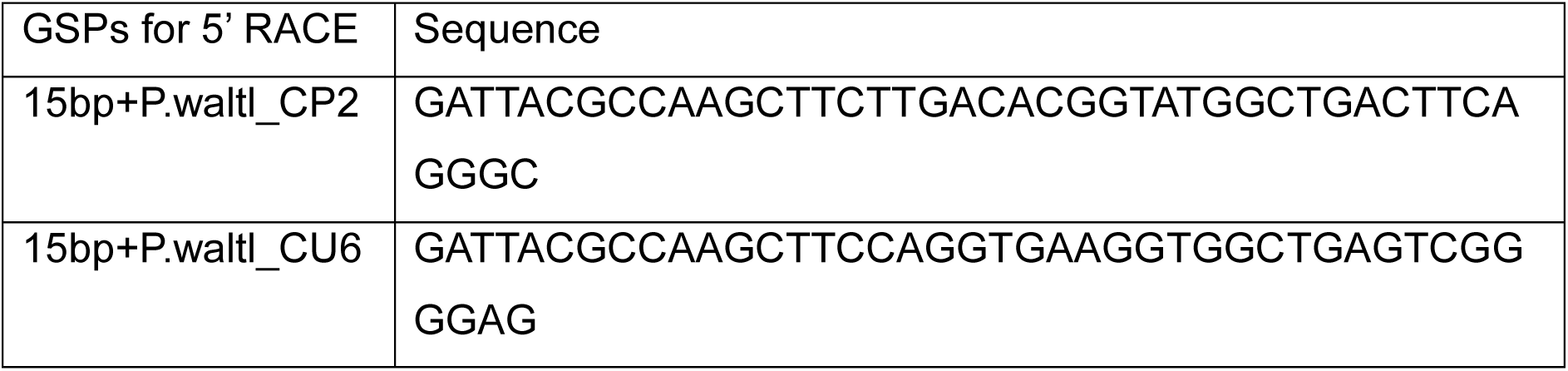
Primers for 5’ RACE.

### Skin grafting

Donor animal was narcotized in 0.1% Tricaine and transferred onto paper towels moistened with 0.05% Tricaine for the procedures. A defined full-thickness skin region (Ø 3mm diameter) was excised from either dorsal side near the base of the tail. Excised skin flaps were maintained briefly in amphibian PBS (APBS) with Ca^2+^ and Mg^2+^. Recipient animals were narcotized in 0.1% Tricaine and also transferred onto paper towels moistened with 0.05% Tricaine for the procedure. The graft bed was prepared by excising a skin flap of identical size at the dorsal base of the trunk (midpoint above the hindlimbs) or on the left forelimb. The donor skin was transferred and patted onto the recipient bed immediately after the graft bed was prepared. Grafted skin was held in place with VetBond™ (butyl cyanoacrylate) which also minimizes desiccation before recipient animal is reintroduced to water. For skin transplant into the blastema, a quarter of the excised skin is folded into the medium bud blastema without applying VetBond™. Recipient animals remained for two hours on paper towels moistened with 0.05% Tricaine before reintroduction to clean shallow conditioned salamander water at room temperature, which was replaced daily for 7 days (with slight water level increment from the fourth day) before transferring to deeper water. Animals were monitored daily for graft adherence and maintained individually to prevent cannibal attacks on grafts. Animals were imaged every 7-10 days for signs of graft rejection.

### Immunofluorescent staining and imaging

Spleen samples were harvested at the post-metamorphic stage from *P. waltl Rag1*^+/−^, *Rag1*^−/−^, and wild-type animals. Spleens were immediately fixed in 4% paraformaldehyde (PFA) (ThermoFisher 043368.9M) containing 0.1% Triton X-100 in DPBS, followed by cryoprotection in a sucrose gradient (10%, 20%, and 30% in DPBS). Samples were embedded and sectioned at a thickness of 12 μm. Sections were stained with antibodies against CD3 (Agilent #A045229-2) to label T cells and with phalloidin–488 (ThermoFisher A12379) to visualize F-actin filaments. The anti- CD3 antibody was used at a dilution of 1:100 in DPBS containing 2% normal goat serum (NGS), followed by an anti-rabbit IgG secondary antibody (Cell Signaling Technology Cat# 4414, RRID:AB_10693544) at a dilution of 1:500. Nuclei were counterstained with DAPI (Sigma Aldrich D9542-10MG). Images were acquired using a Leica Thunder imaging system and processed with Leica imaging software.

### Statistics and reproducibility

Sample sizes were tailored uniquely to each experimental aim and the level of experimental blinding. The sample sizes for the F2:Rag1 and F3:Rag1 blinded experiments were based on Mendelian offspring ratio from heterozygous parents, ensuring we have at least, n = 5 per group. Sample sizes for amputation experiments were aimed at n = 6 per group, non-blinded studies without statistical analysis were three – five animals per group. The number of biological and technical replicates are given in the legend for each figure. Heterozygous (*Rag1*^+/-^) and WT (*Rag1*^+/+^) newts have indistinguishable phenotypes before and after all pilot experiments; the number of WT (*Rag1*^+/+^) newts for some experiments reported within were limited to n =1 as reference to the *Rag1*^+/-^ newt control animals or samples. Experiments were repeated at least twice. Statistical analyses performed for each experiment are reported in the figure legends where applicable. Conditions for statistical significance at 95% confidence interval are given in the statistical figure legends where applicable.

## Acknowledgements

We thank all members of the Regenerative Immunology and Systems Immunology labs for their support, feedback, and encouragement. We are grateful to the StemTherapy Strategic Research Area at Lund University for their support. Computations and data storage were enabled by resources provided by LUNARC, The Centre for Scientific and Technical Computing at Lund University.

## Author contributions

Conceptualization: N.D.L.

Methodology: C.H.U., A.G., M.O., N.D.L.

Data curation: E.B.

Investigation: C.H.U., M.O., U.O., A.G., N.L., S.M, C.R.C., N.D.L

Visualization: C.H.U., N.D.L, A.G., C.R.C.

Supervision: N.D.L., A.S., E.E.

Funding acquisition: N.D.L.

Writing—original draft: C.H.U., N.D.L.

Writing—review & editing: All authors

## Data, code, and materials availability

All data are either part of this manuscript or previously published. Previously published single cell RNA seq data are available at Sequence Read Archive SRP167700. All code is available at: https://github.com/RegenImm-Lab/adaptive_immune_limb_regen

