## Supplementary material for "Adaptive immunity is dispensable for salamander appendage regeneration": Supplmental Figures

### Supplemental Figure 1

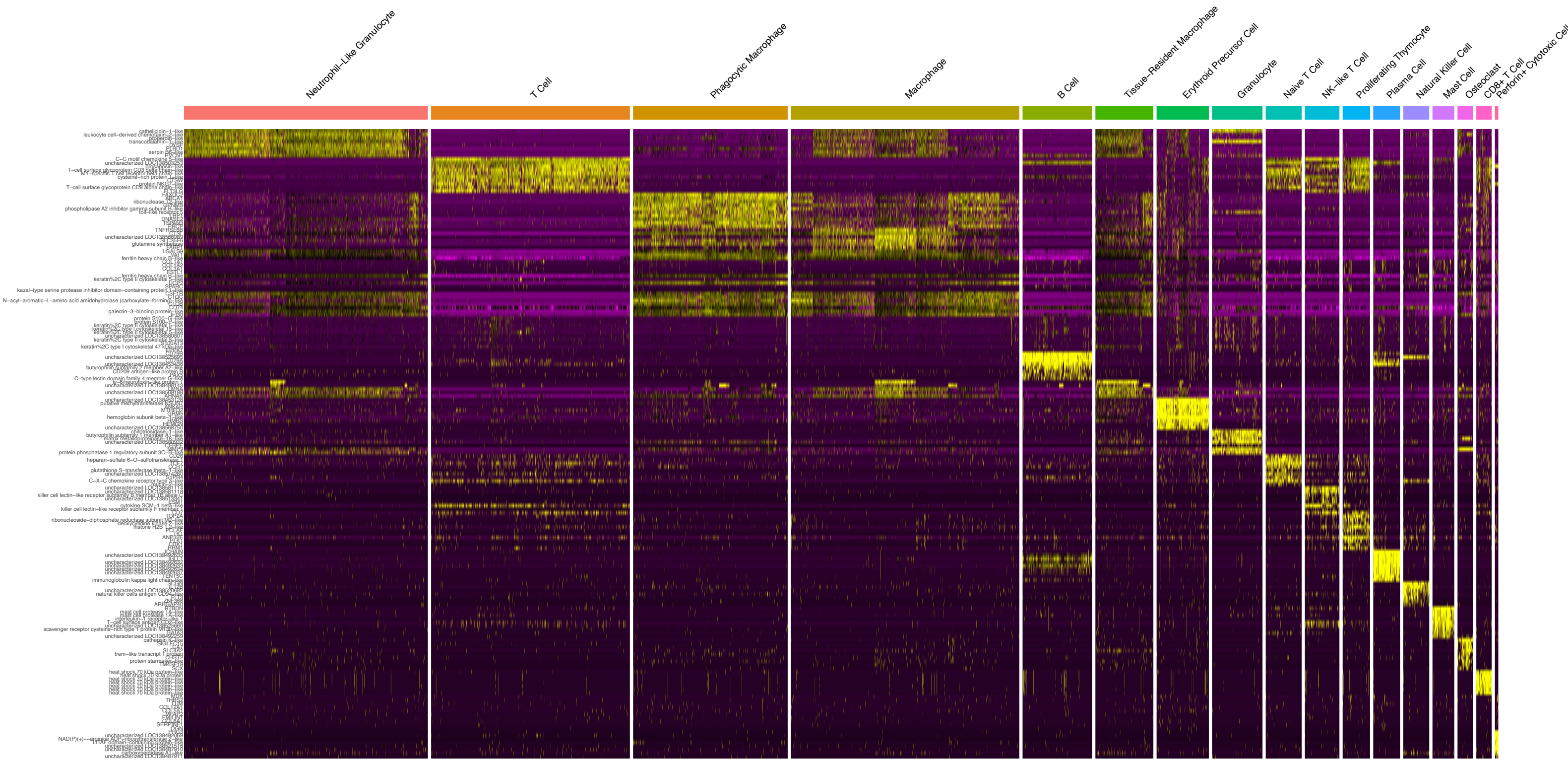


**Supplemental Figure 1: Heatmap showing top 10 markers for each cell cluster.** If a gene name was available it was maintained for each row. If the gene was a LOC identifier it was replaced with the gene annotation information, when available. Genes were filtered for logFC > 2.

### Supplemental Figure 2

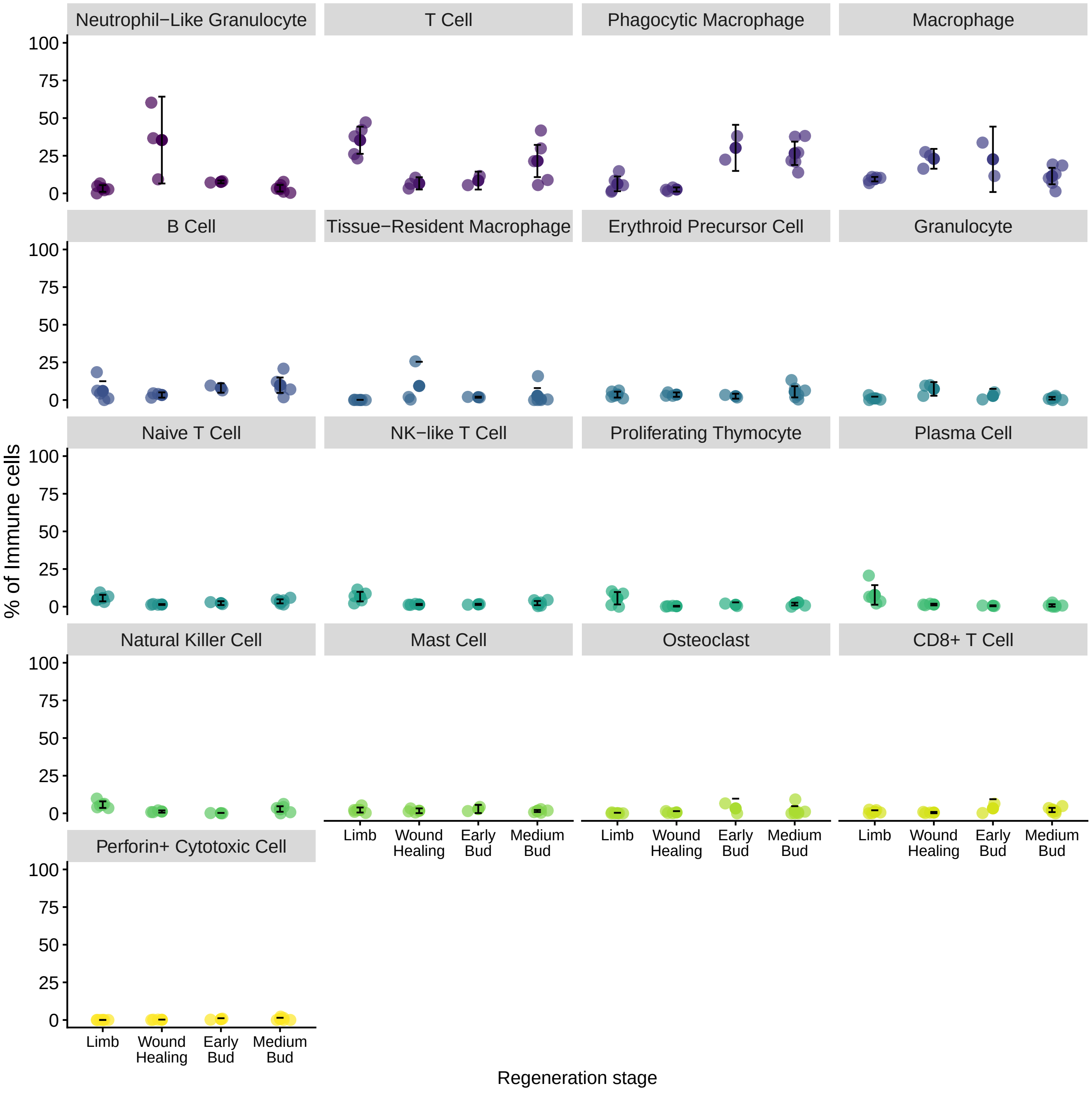


**Supplemental Figure 2: Immune populations are dynamic across regeneration.** Dot plot visualizing the percentage of total immune cells of each annotated immune cell type. Each dot represents a biological replicate.

### Supplemental Figure 3

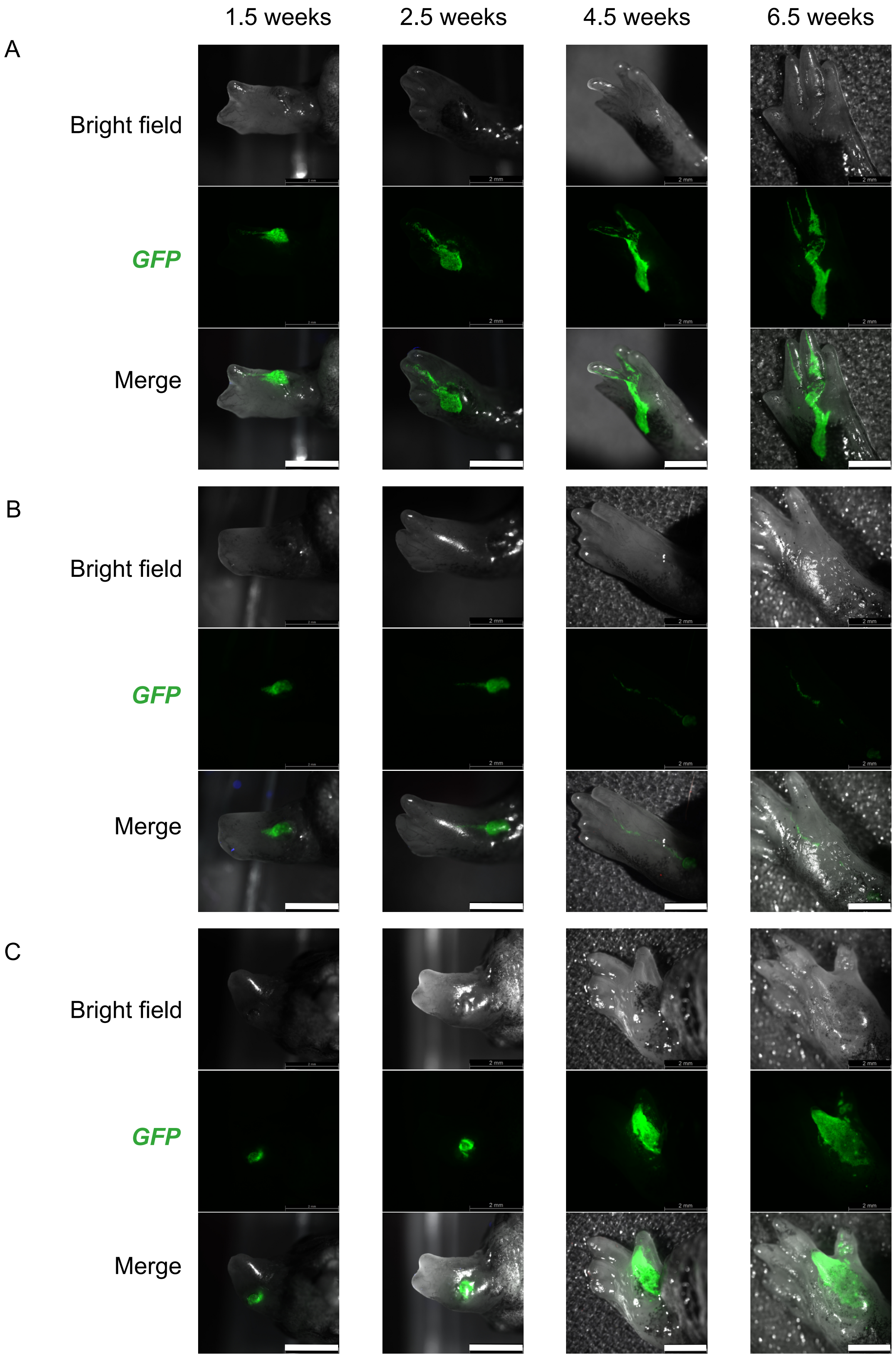


**Supplemental Figure 3: Exemplary images showing fates of allograft GFP skin tissues in three animals represented by panels A, B and C.** Allograft skin contributed to (A) parts of the hand and digits, (B) blood vessels and (C) hand and supernumerary digit. Scale bars = 2mm.

### Supplemental Figure 4

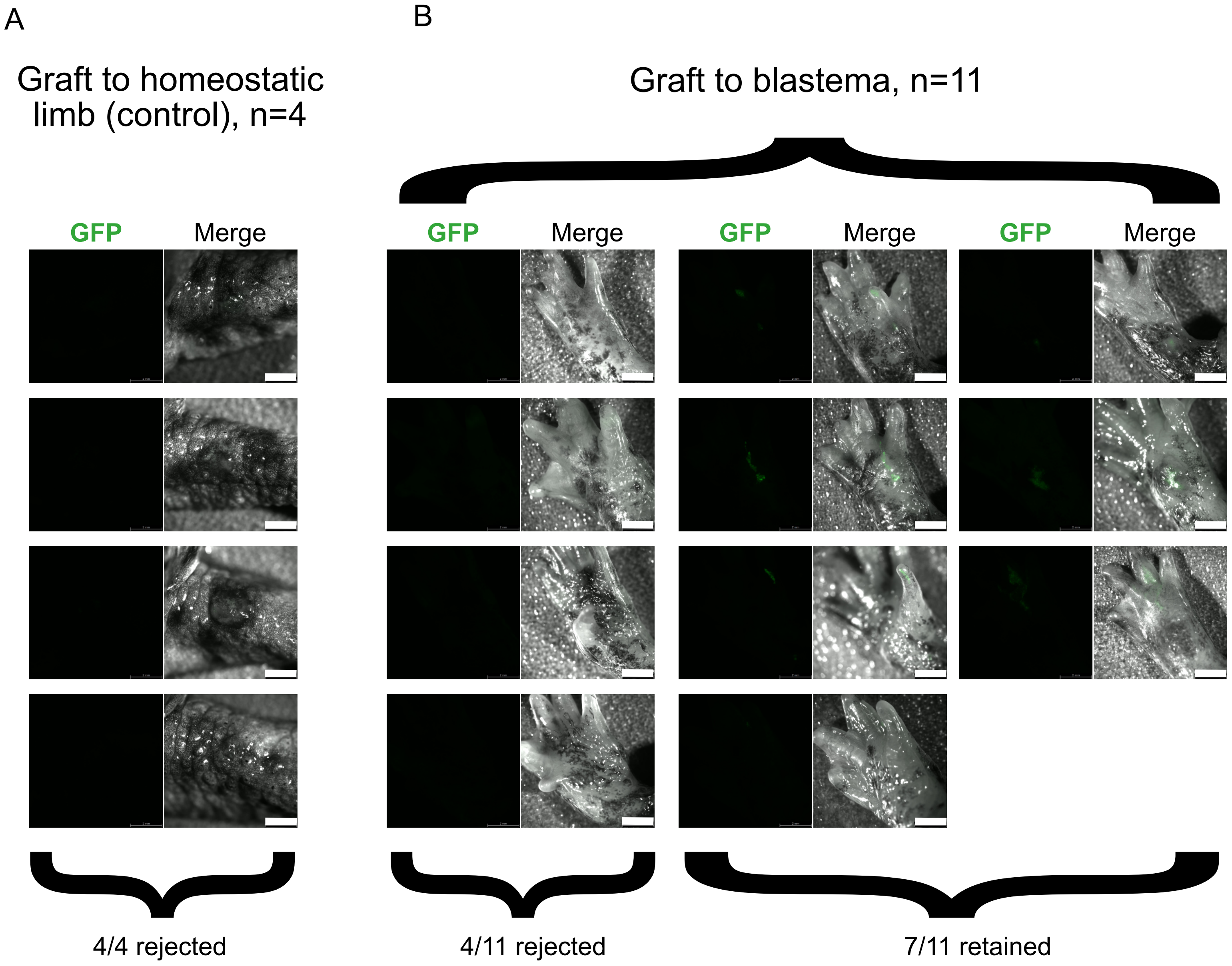


**Supplemental Figure 4: Images depicting graft fates at the final time point of 11.5 weeks.** (A) 4 out of 4 control animals rejected the GFP skin allograft. (B) 4 out of 11 animals grafted in the blastema rejected GFP skin allograft, while 7 out of 11 retained GFP skin allograft at the same time point. Scale bars = 2mm.

### Supplemental Figure 5

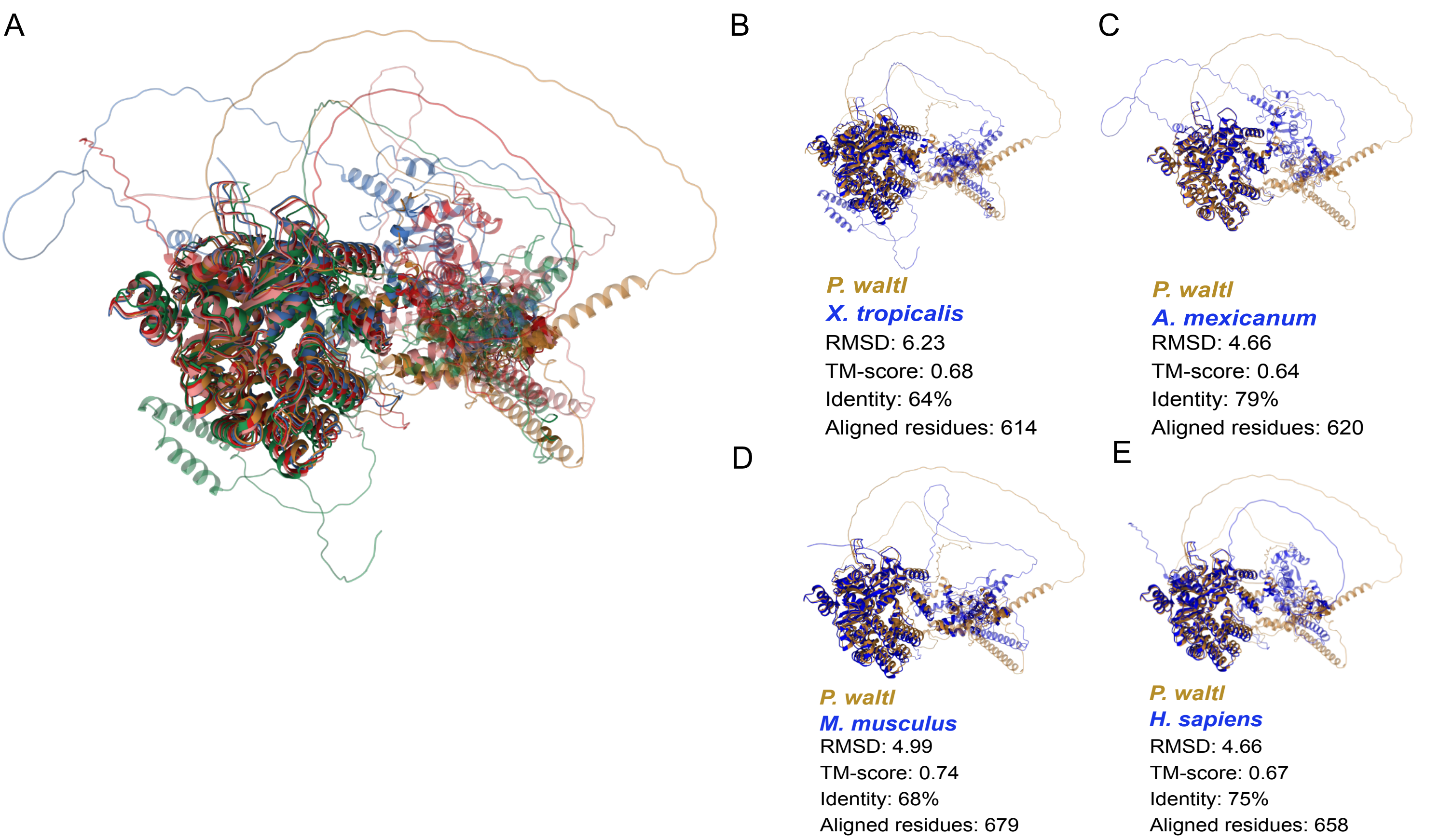


**Supplemental Figure 5: Comparing predicted Rag1 structure across species.**
**(A)** Predicted 3D conformation of *P. waltl* Rag1 protein superimposed on predicted Rag1 protein 3D structures of orthologs. **(B-E)** Superimposition of predicted *P. waltl* Rag1 protein 3D structure individually on compared orthologs **(B)** *X. tropicalis*, **(C)** *A. mexicanum*, **(D)** *M. musculus* **(E)** *H. sapiens*. Predictions were generated using Foldseek2.

### Supplemental Figure 6

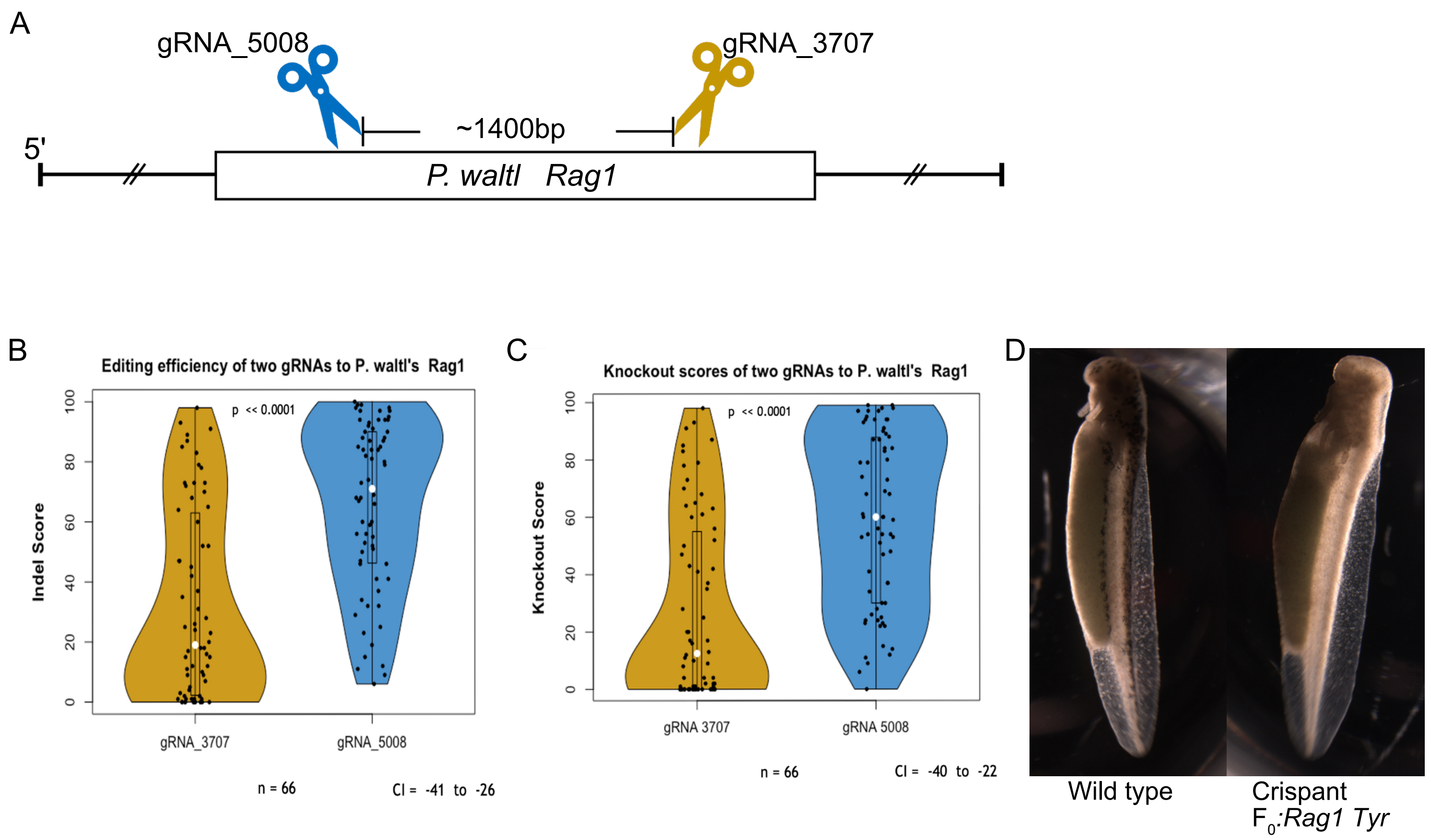


**Supplemental Figure 6: Establishing *Rag1* deficient salamanders.**

**(A)** Target locations of gRNAs on *P.waltl Rag1*. Scissors indicate the gRNA-Cas9 cut sites.**(B-C)** Violin plots showing differences in the median **(B)** editing efficiencies and **(C)** knockout scores of the two *Rag1* targeting gRNAs used. Black dots represent individual animal, n = 66, $p < 0.0001$

**(D)** Loss of pigmentation observed at 7 days post-CRISPR/Cas9 in *Tyrosinase*-targeting gRNA injected embryos.

### Supplemental Figure 7

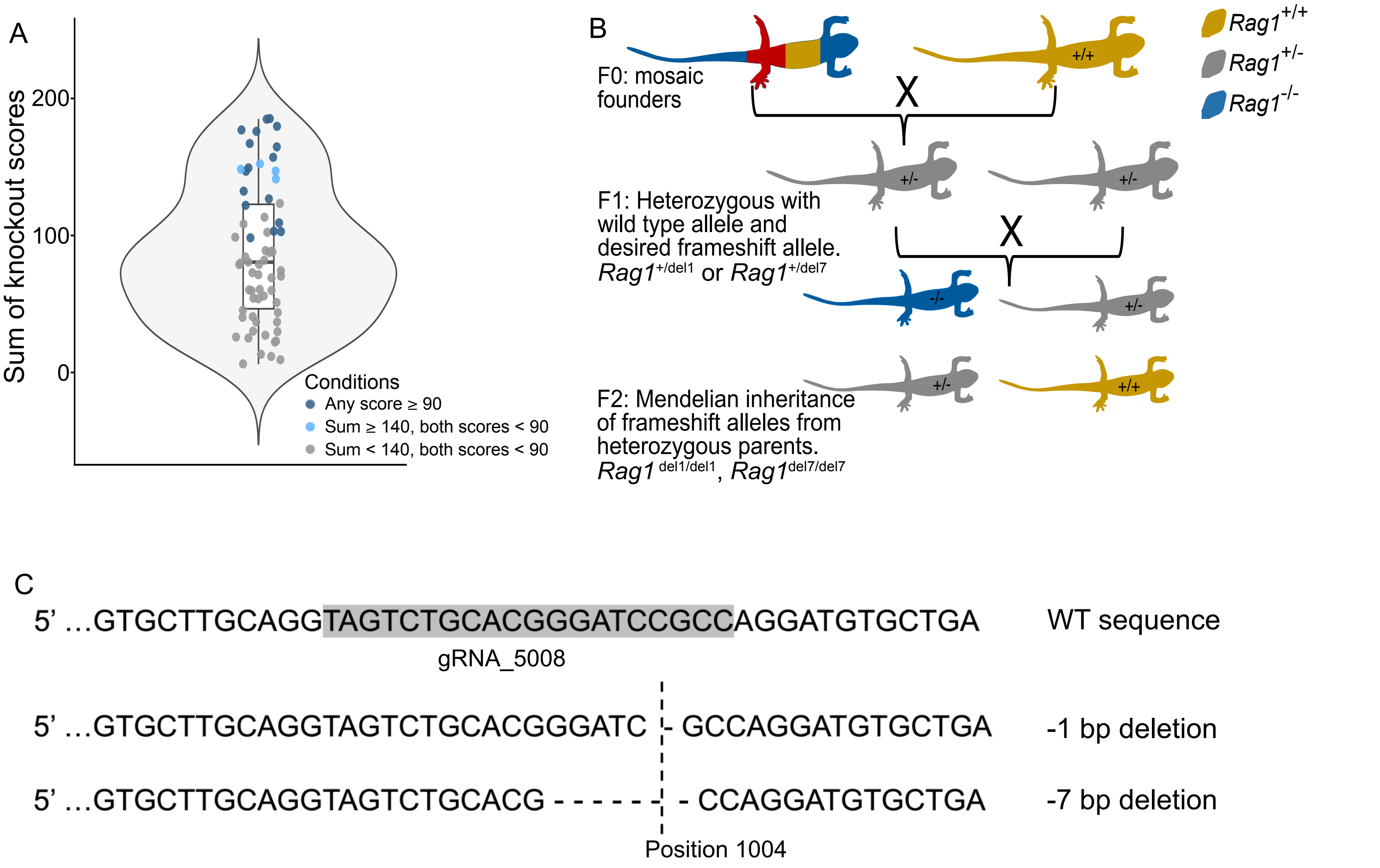


**Supplemental Figure 7: Breeding and resulting deletions observed in the two *Rag1*-/- newts lines.** **(A)** Sum of the knockout scores on the *Rag1* gene for each $F_0$:*Rag1*-crispant salamander, n = 66. Blue (light and dark) data points dedicated to "*Rag1* High edit" in Figure 5A. **(B)** Mendelian Scheme for generating mutant $F_2$:*Rag1* animals from $F_{0:}$crispant founders. Crispant founders carrying the desired mutations in the *Rag1* gene were outcrossed with WT salamanders, and the resulting offspring carrying one mutant allele were in-crossed to produce third generation offspring having both alleles either mutant (*Rag1*$^{-/-}$) or heterozygous (*Rag1*$^{+/-}$) or WT (*Rag1*$^{+/+}$) in a Mendelian distribution. (C) Locus of CRISPR cut site on the newt *Rag1* gene, yielding two mutations (*Rag1*$^{del1/del1}$ and *Rag1*$^{del7/del7}$) for the *Rag1*-/- newts used in the study.

### Supplemental Figure 8

A

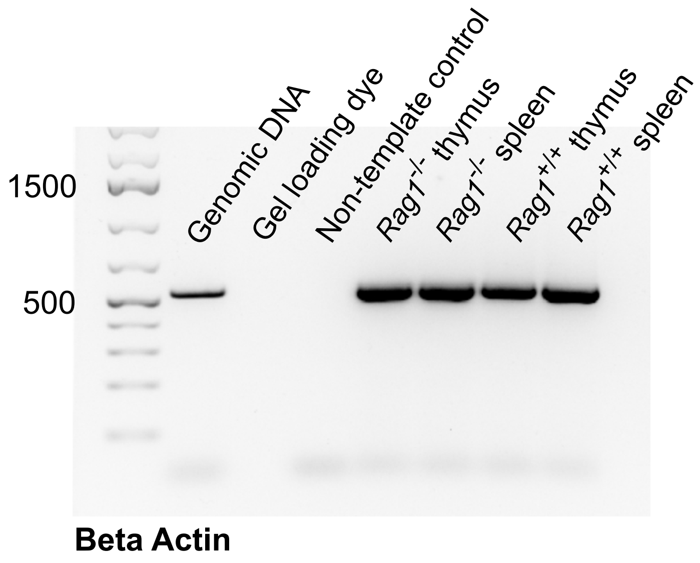


B

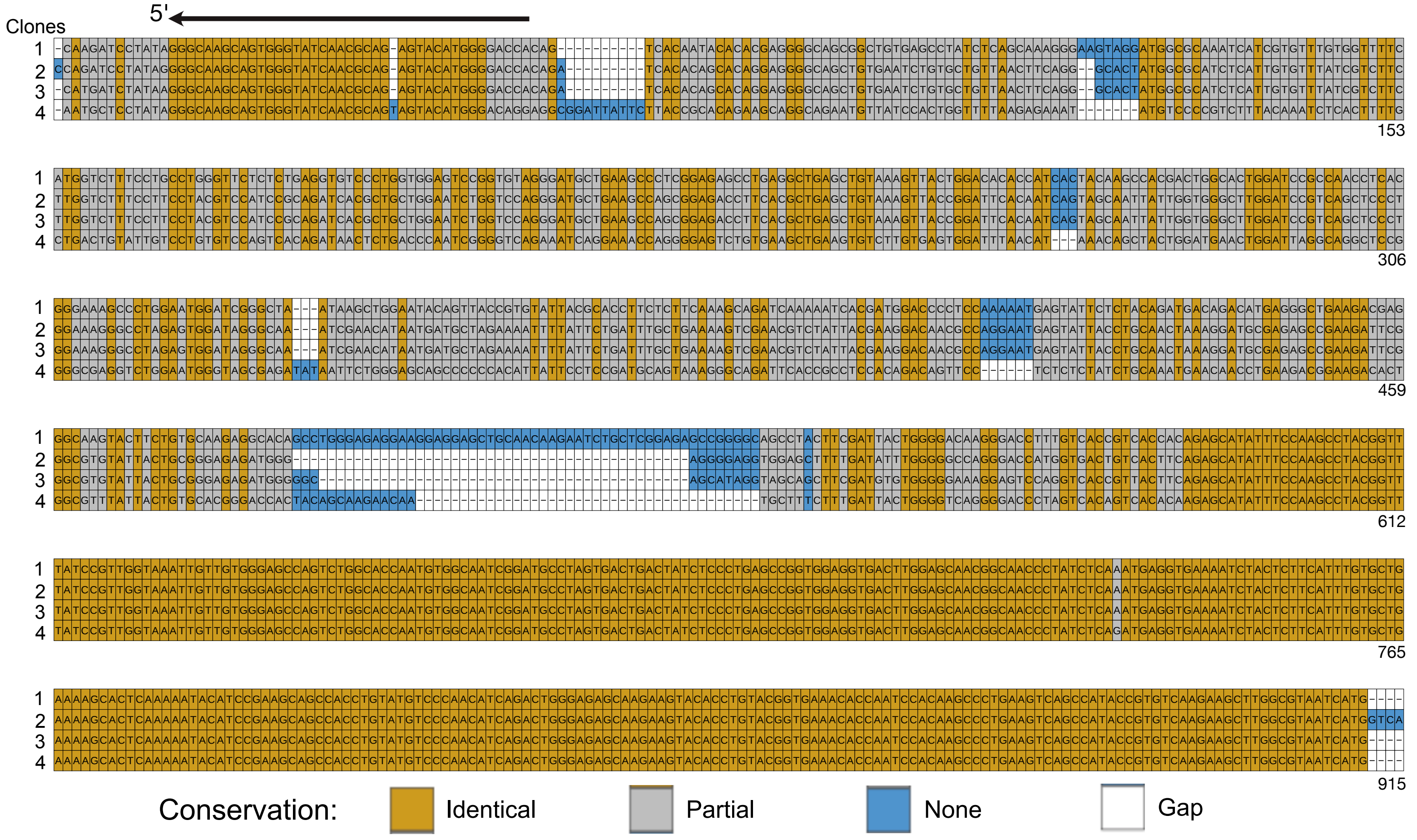


**Supplemental Figure 8: Verification of RACE PCR amplification of antigen receptors.** **(A)** PCR for beta actin as quality control for template starting RACE cDNA across samples and controls. **(B)** Alignment of sequences from four randomly selected clones of the *Rag1*$^{+/+}$ RACE products showing variability and conservation of the clones: identical (orange), partial (grey), no similarity (blue), gap (white). While, the 3' regions show identical conservation, the middle regions up to the 5' regions show variability in the sequences. Top arrow points to the 5' direction
